# Deep proteomic analysis of microglia reveals fundamental biological differences between model systems

**DOI:** 10.1101/2022.07.07.498804

**Authors:** Amy F Lloyd, Anna Martinez-Muriana, Emma Davis, Michael JD Daniels, Pengfei Hou, Renzo Mancuso, Alejandro J Brenes, Ivana Geric, An Snellinx, Katleen Craessaerts, Tom Theys, Mark Fiers, Bart De Strooper, Andrew JM Howden

**Affiliations:** Cell Signalling and Immunology, University of Dundee, Dundee, UK; VIB Centre for Brain & Disease Research, Leuven, Belgium; Department of Neurosciences, Leuven Brain Institute, KU Leuven, Leuven, Belgium; The Francis Crick Institute, London, UK; UK Dementia Research Institute at UCL, University College London, London, UK; UK Dementia Research Institute at University of Edinburgh, Edinburgh, UK; Department of Biomedical Sciences, University of Antwerp, Antwerp, Belgium; MIND lab, VIB Centre for Molecular Neurology, VIB, Antwerp, Belgium; Centre for Gene Regulation and Expression, University of Dundee, Dundee, UK; Department of Neurosciences, Research Group Experimental Neurosurgery and Neuroanatomy, KU Leuven, Leuven, Belgium

## Abstract

Using high resolution quantitative mass spectrometry, we have generated the most comprehensive human and mouse microglia proteomic datasets to date, consisting of over 11,000 proteins across all six microglia groups. Microglia from different sources share a core protein signature of over 5600 proteins, yet fundamental differences are observed between species and culture conditions, indicating limitations for human disease modelling in mouse or in *in vitro* cultures of microglia. Mouse *ex vivo* microglia show important differences at the proteome level such as differential expression of inflammation and Alzheimer’s Disease associated proteins. We identify a tenfold difference in the protein content of *ex vivo* and *in vitro* cells and significant proteome differences associated with protein synthesis, metabolism, microglia marker expression and environmental sensors. Culturing microglia induces rapidly increased growth, protein content and inflammatory protein expression. These changes can be restored by engrafting *in vitro* cells into the brain, with xenografted hESC-derived microglia closely resembling microglia from human brain. This data provides an important resource for the field and highlights important considerations needed when using model systems to study human physiology and pathology of microglia.

## Introduction

Microglia, the resident immune cells of the central nervous system (CNS) are the first responders to pathogenic invasion and damage. Developmentally, microglia play critical roles in axon guidance [1, 2], synaptic pruning [3, 4], supporting neuronal survival [5, 6] and myelin maintenance [7, 8]. Although post-development, microglia are maintained in a homeostatic state of constant surveillance of their environment, they rapidly activate upon stimulation, adopting a plethora of phenotypes that are mainly defined at the gene expression level. Depending on the stimulus, microglia can range from activating to a potent inflammatory state, to that of a tissue-reparative, inflammatory-resolving state [9, 10]. Sustained activation of the former is a pathological hallmark of neurodegenerative diseases such as Alzheimer’s disease [11, 12] and multiple sclerosis [9, 10], manifesting a chronic inflammatory environment and progressive neuronal and glial cell death.

Manipulation of *in vitro*, *ex vivo* and *in vivo* models has been critical in our understanding of microglia biology and function, however the limitations of these models to our understanding of human diseases is unknown. Transcriptomic studies have uncovered the heterogenic landscape of murine and human microglia throughout the CNS in health and disease [13–16], yet disparity between the transcriptome and proteome [17–21] means that much is still to be known about microglia. Pearson correlation coefficients between protein and gene expression in a variety of CNS cell types, including microglia, range from 0.40-0.45, highlighting the unreliability of the transcript to predict protein expression [19]. Single cell transcriptomics yield significantly lower identified genes than proteomics, which can also cause underrepresentation of cellular responses [20]. The impact of this has been powerfully demonstrated in Alzheimer’s Disease (AD) brain tissue where around half of all protein modules identified were not seen at the transcript level [17]. Two of these modules were disease-specific, with increased expression of the MAPK module correlating to cognitive decline, highlighting the importance of defining disease kinetics at the protein level. Post-transcriptional and translational alterations, and varying rates of protein degradation have also been blamed for this transcript-proteome disconnect [18, 22] and therefore make it difficult to interpret gene expression data biologically and functionally.

Proteins are the main mechanistic mediators of cell phenotypes and function, yet the human microglia proteome remains poorly defined. Unravelling the human microglia proteome and understanding how well experimental systems effectively model the biology of human microglia is of great importance both experimentally and clinically. Indeed, differences in human and mouse microglia gene and protein expression in neurodegeneration have been documented [15, 20, 23, 24], with clinical relevance demonstrating the disease associated microglia (DAM) signature, initially defined in mouse models of amyloidosis, is incomplete in human microglia [20]. *In vitro* microglia allow greater pharmacological and temporal manipulation to model cellular and molecular responses, and although differential responses to inflammatory stimuli have been documented in mouse microglia *in vitro* compared to *in vivo* [25, 26], the full extent of their relevance, particularly at the protein level, is unknown. Therefore, a detailed proteomic map of human and mouse microglia from *in vitro* and *ex vivo* sources is warranted to fully understand the advantages and limitations of experimental systems used to model human microglia in disease contexts.

To investigate the similarities and differences between microglia model systems, we used high resolution quantitative mass spectrometry to map the proteomes of *ex vivo* human microglia, *ex vivo* and *in vitro* cultured mouse microglia, *in vitro* cultured and xenografted human embryonic stem cell (hESC)-derived microglia and the BV2 cell line. We have identified and quantified in total over 11,000 different microglia proteins with deep coverage across all microglia groups, generating the most comprehensive microglia proteomic resource to date. We reveal striking differences between *in vitro* and *ex vivo* cells and their capacity for protein synthesis, energy production and their ability to sense and respond to their environment. This highly plastic reprogramming is observed by comparing microglia transitions both from *ex vivo* to *in vitro* (mouse microglia protein signatures sorted from the brain compared to 7 days in culture), and vice-versa (human ESC-microglia *in vitro* and after engraftment into the mouse brain).

This data provides a valuable resource to the field which can be easily interrogated using our web portal https://data.bdslab.org/Lloyd2024 (*trial version available until publication*). Furthermore, this resource details fundamental new insights into the biology of microglia models and will help better understand the role of these cells in health and disease.

## Results

### Proteomics reveals vast differences between in vitro and ex vivo microglia

Quantitative mass spectrometry was used to uncover proteomes of microglia from human surgical donors, human H9 ESC-derived microglia from *in vitro* cultures and from mouse brain xenografts, and *in vitro* and *ex vivo* adult mouse microglia. (Fig. 1a). For a full comparison of models commonly used for understanding microglia biology we also analysed the proteome of the BV2 cell line (Supplemental Fig. 1, 2). All sample information can be found in Supplementary File 1. Full proteome data for all samples include BV2 cells is also available in Supplementary File 1.

**Figure 1.**
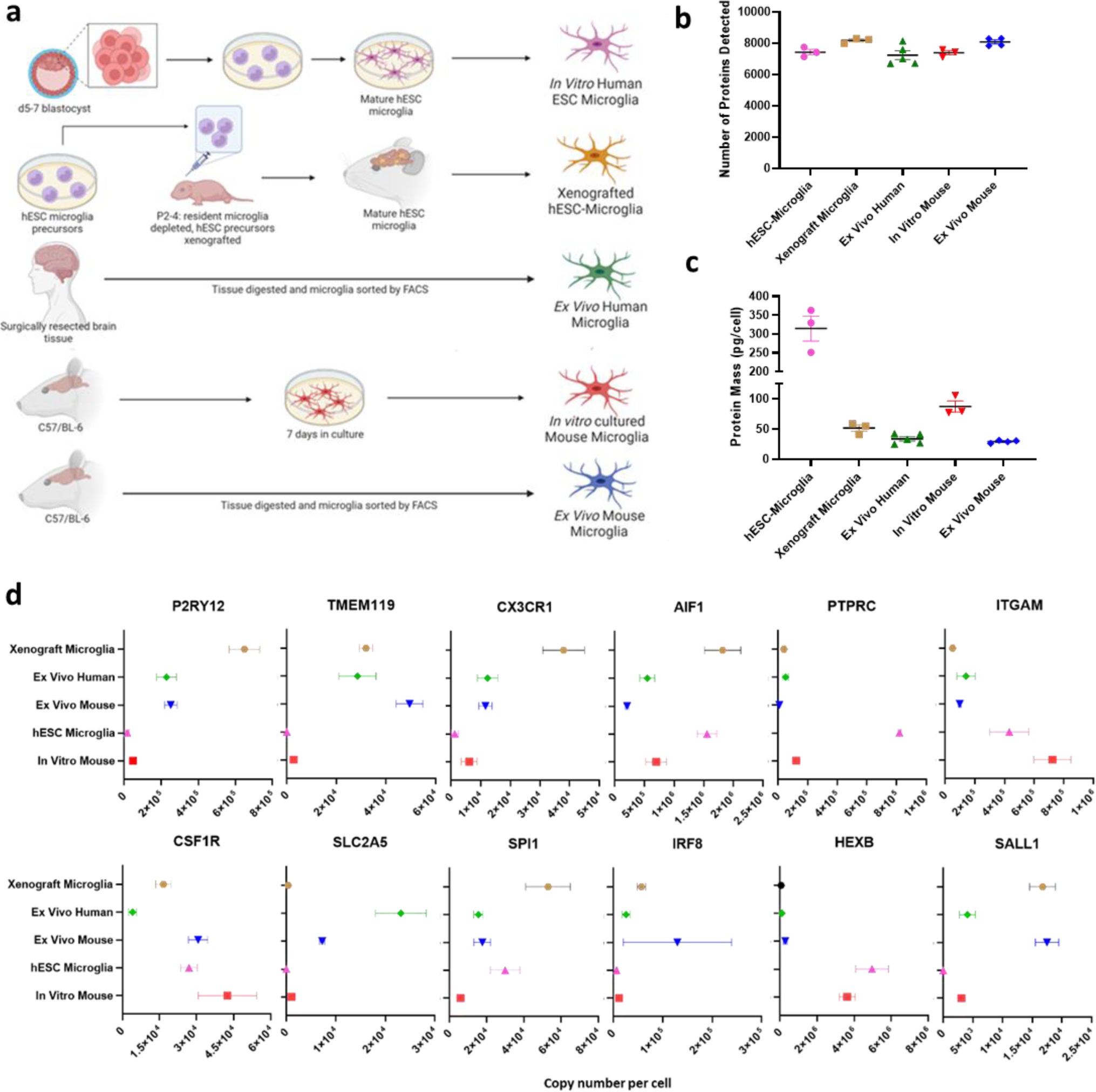
a) Schematic of microglia collection methods and sources. *In vitro* hESC-microglia are derived from the H9 ESC line (n=3). Xenograft microglia are H9 hESC-derived microglia isolated from mice aged 3 months (n=3). *Ex vivo* human microglia (n=5). *In vitro* mouse microglia (n=3). *Ex vivo* mouse microglia (n=4). **b)** Total number of proteins detected in each microglia subset ±SEM. **c)** Protein mass of each microglia subset, measured in picograms (pg) per cell ±SEM. **d)** Quantified protein expression (copy numbers per cell) of core microglia proteins (mean copy number ±SEM).

*Ex vivo* microglia were isolated from human and mouse brains by fluorescence-activated cell sorting (FACS) (Supplemental Fig. 1a, b). Human microglia were isolated from surrounding tissue, identified as normal by pathological inspection, of surgically resected temporal lobe tissue of epilepsy patients (Supplemental Fig. 1c), as previously described [13, 15, 27-29]. To generate a deep microglia proteome for each subset, samples were fractionated by high-performance liquid chromatography (HPLC) into 16 fractions prior to mass spec analysis, enabling a more in-depth proteome coverage that includes the detection of less abundant proteins, as well as higher accuracy of quantification. It is important to note that if a protein was not detected in a sample, we cannot conclude that the protein was absent. Failure to detect a protein can be the result of a protein being expressed below the threshold of detection or due to physiochemical properties of the peptides impacting their ionisation efficiency. Nevertheless, in total, 9437 human and 9633 mouse proteins were identified (Supplementary File 1). The average numbers of proteins identified across bio-replicates for each subset varied only slightly, ranging between 7234 and 8315 for all samples (Fig. 1b, Supplemental Fig. 1d, e). Of these proteins, a core proteome of 5685 proteins was identified across all six microglia groups (Supplemental Fig. 1f). Gene ontology identified core proteome networks associated with immune responses, type 1 interferon signalling and protein targeting to lysosomes (Supplemental Fig. 1g).

Immediate striking differences between samples were noted in their total protein mass; *in vitro* cultured hESC-microglia contained ∼300pg of protein per cell, around 10-fold greater than *ex vivo* human and mouse microglia (Fig. 1c, Supplemental Fig. 1h). Interestingly, this correlated with decreased inhibition of the eukaryotic initiation factor 4 (eIF4) complex as regulated by the translational repressor programmed cell death 4 (PDCD4) (Supplemental Fig. 2a). As one molecule of PDCD4 binds two molecules of eIF4A1 [30], we calculated the ratio of PDCD4:eIF4A1 for each microglia group. The tightest regulation of translation as shown by the lowest numbers of eIF4A molecules per PDCD4 molecule, was observed in xenograft and *ex vivo* microglia (Supplemental Fig. 2b). Histone content across samples was comparable (Supplemental Fig. 2c).

Comparison of expression of core microglia proteins revealed large variations across microglia groups (Fig. 1d, Supplemental Fig. 2d). We noted that expression of homeostasis markers P2RY12, TMEM119 and CX3CR1 were high in *ex vivo* groups, but low or undetected in *in vitro* groups. Overall expression of most microglia markers was notably lower in *in vitro* cells (Fig. 1d, Supplemental Fig. 2d). Principal Component Analysis (PCA) of protein copy number expression in all microglia groups prior to differential analysis revealed that environment (*in vitro* vs *ex vivo*; PC1) and species (mouse v human; PC2) are the major drivers of variance between samples (Supplemental Fig. 2e). Although our microglia proteomes are derived from both male and female samples, we do not find that these differences in microglia protein signatures are driven by sex (Supplemental Fig. 2f).

Weighted correlation network analysis (WGCNA) further highlighted the proteomic divergence between *in vitro* and *ex vivo* microglia (Fig. 2a, b, Supplemental File 1). Protein modules were annotated using gene ontology enrichment analysis and assigned individual colours. Proteins in the N/A module were not significantly associated with a GO term, however, all proteins assigned to each module can be further interrogated in Supplemental File 1. Protein modules associated with cell growth and metabolism and translation were among the most strongly enriched in *in vitro* microglia, with RNA processing and actin regulatory complex Arp2/3 enriched in *ex vivo* microglia. Interestingly, species-specific protein modules were also identified, including enrichment of fatty acid beta oxidation in mice, and RNA splicing, mitochondrial translation elongation and mTOR signalling enriched in human (Fig. 2a, b). Pearson correlations between weighted protein correlation network modules identified meta modules with top ranking proteins in each module displayed (Fig. 2b). Modules clustered into 5 groups based on enrichment in either *in vitro* or *ex vivo* groups, and human or mouse. 2 clusters showed enrichment in *ex vivo* and mouse microglia groups. These two clusters included strong correlation of protein modules associated with translation, cell growth and metabolism and fatty acid oxidation, fibrinolysis, synapse pruning and astrocyte development.

**Figure 2.**
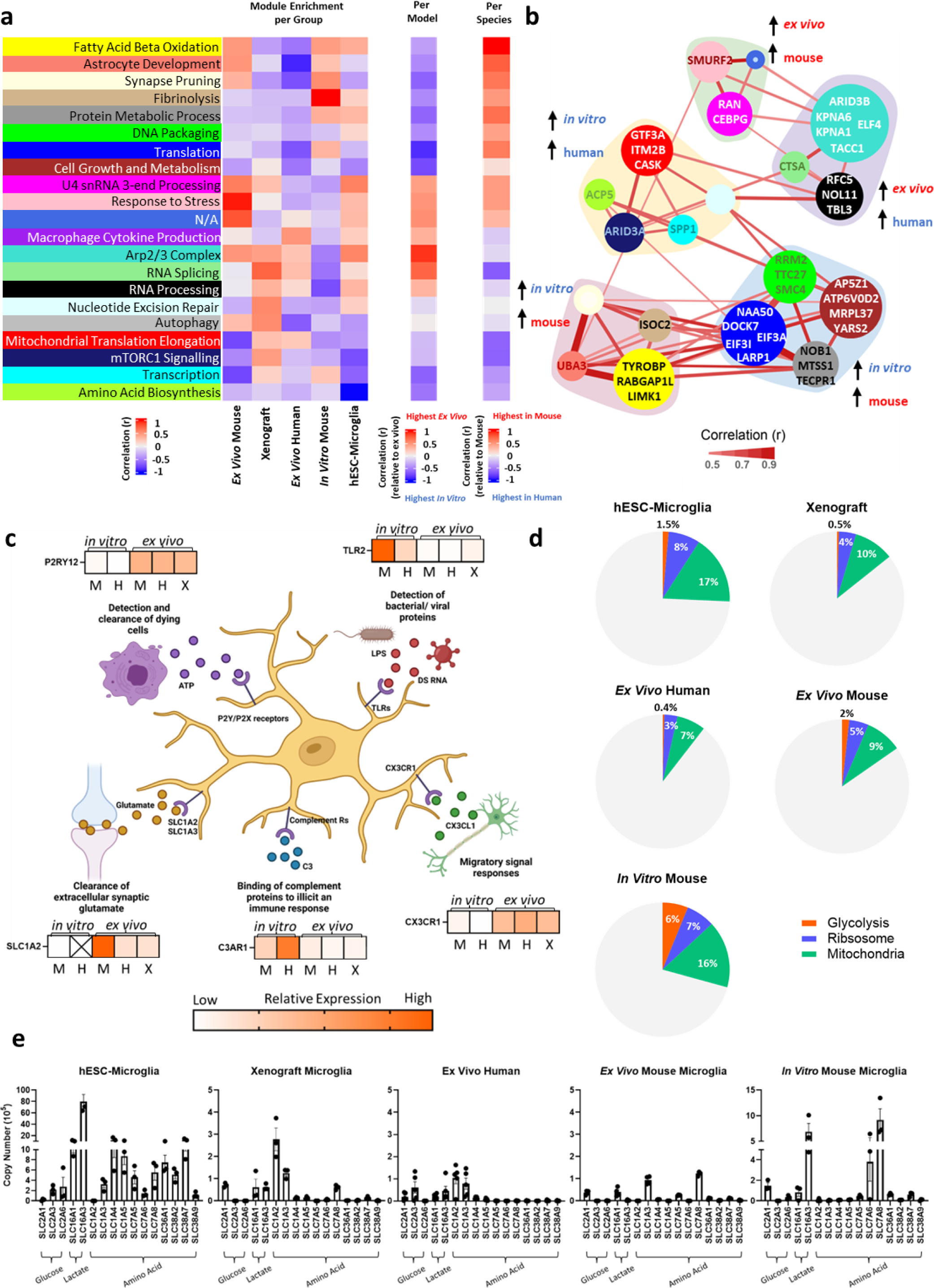
a) Heatmap representing weighted protein correlation network analysis (WGCNA) used to determine enriched protein modules. Each protein module is assigned a colour. Module enrichment per group highlights relative expression of protein modules for each microglia group using Pearson Correlation Coefficient (r): red = relative enrichment of protein modules, blue = relative downregulation of protein modules. Module enrichment per model maps protein modules correlated (Pearson, r) to either *ex vivo* (red) or *in vitro* (blue) microglia groups. Strong red/blue indicates strong correlation of protein modules associated with model. Module enrichment per species maps protein modules correlated (Pearson, r) to either mouse (red) or human (blue). Strong red/blue indicates strong correlation of protein modules associated with species. **b)** Pearson correlations between weighted protein correlation network modules grouped into meta modules defined by hierarchical clustering. Colours correspond to module names as indicated in Fig. 1a. Proteins displayed in each module are the top-ranking proteins in each module, ranked by module membership (where module membership is at least >0.5). Module node size scales with the number of proteins within each module. As meta-modules largely cluster based on species and model, this trait correlation is labelled for each meta-module. **c)** Schematic of different facets of the microglia sensome with key receptors and heatmaps of relative copy number expression between microglia groups from low (white) to high (orange). Order of samples: M: *in vitro* mouse (n=3), H: *in vitro* hESC-microglia (n=3), M: *ex vivo* mouse (n=4), H: *ex vivo* human (n=5) and X: xenograft (hESC-microglia; n=3). **d)** Pie charts displaying the proportion of total protein mass of each microglia group associated with glycolysis (orange), ribosome (blue) and mitochondria (green). **e)** Total copies per cell (displayed as log10 copy numbers) of glucose, lactate and amino acid nutrient transporters for hESC-microglia (n=3), xenograft microglia (n=3), *ex vivo* human microglia (n=5), *ex vivo* mouse microglia (n=4) and *in vitro* mouse microglia (n=3) ±SEM.

Relative expression of sensome-associated microglia proteins could also be separated between *in vitro* and *ex vivo* microglia (Fig. 2c, Supplemental Fig. 2g). As well as P2RY12, a chemoreceptor responsible for the detection of ATP released from synaptic transmission and dying cells, and CX3CR1, important for responding to migratory signals released from neurons, SLC1A2 a nutrient transporter involved in the detection of extracellular synaptic glutamate, was also predominantly expressed in *ex vivo* microglia. Conversely, complement 3 (C3) receptor C3AR1 and toll-like receptor (TLR) 2, associated with inflammatory responses, were enriched in *in vitro* microglia.

As the *in vitro* environment exposes cells to high levels of growth factors, serum and cytokines that can all greatly impact function and phenotype [31] we sought to understand how the *in vitro* and *ex vivo* environments manifest differences in bioenergetics. We uncovered the proportions of the total protein mass for each sample dedicated to glycolysis, ribosome, and mitochondrial proteins (Fig. 2d) and found that *in vitro* cells showed increases in all three compared to their species-specific *ex vivo* counterparts. Interestingly, *ex vivo* mouse microglia had a higher proportion of their protein mass dedicated to glycolysis compared to all human *ex vivo* and *in vitro* groups, however this was tripled in *in vitro* mouse microglia, suggesting a higher basal glycolytic rate in mouse microglia compared to human. Expression of nutrient transporters differed greatly across microglia groups (Fig. 2e) with lactate and amino acid transporters highly expressed in *in vitro* microglia. *In vitro* hESC-microglia in particular showed strikingly high expression of a wide range of amino acid transporters compared to all other groups. Overall, this data highlights the impact of the *in vitro* environment and increased nutrient exposure on microglia bioenergetics which may be a major influence on phenotypes.

### Fundamental differences in protein expression patterns of ex vivo mouse and human microglia

We sought to understand and define the *ex vivo* mouse and human derived microglia proteomes and compare their composition (Fig. 3a). Although variation between human samples is expected, the number of proteins detected (Supplemental Fig. 3a) and protein mass (Supplemental Fig. 3b) were similar across all human samples, with significant overlap of proteins expressed (Supplemental Fig. 3c).

**Figure 3.**
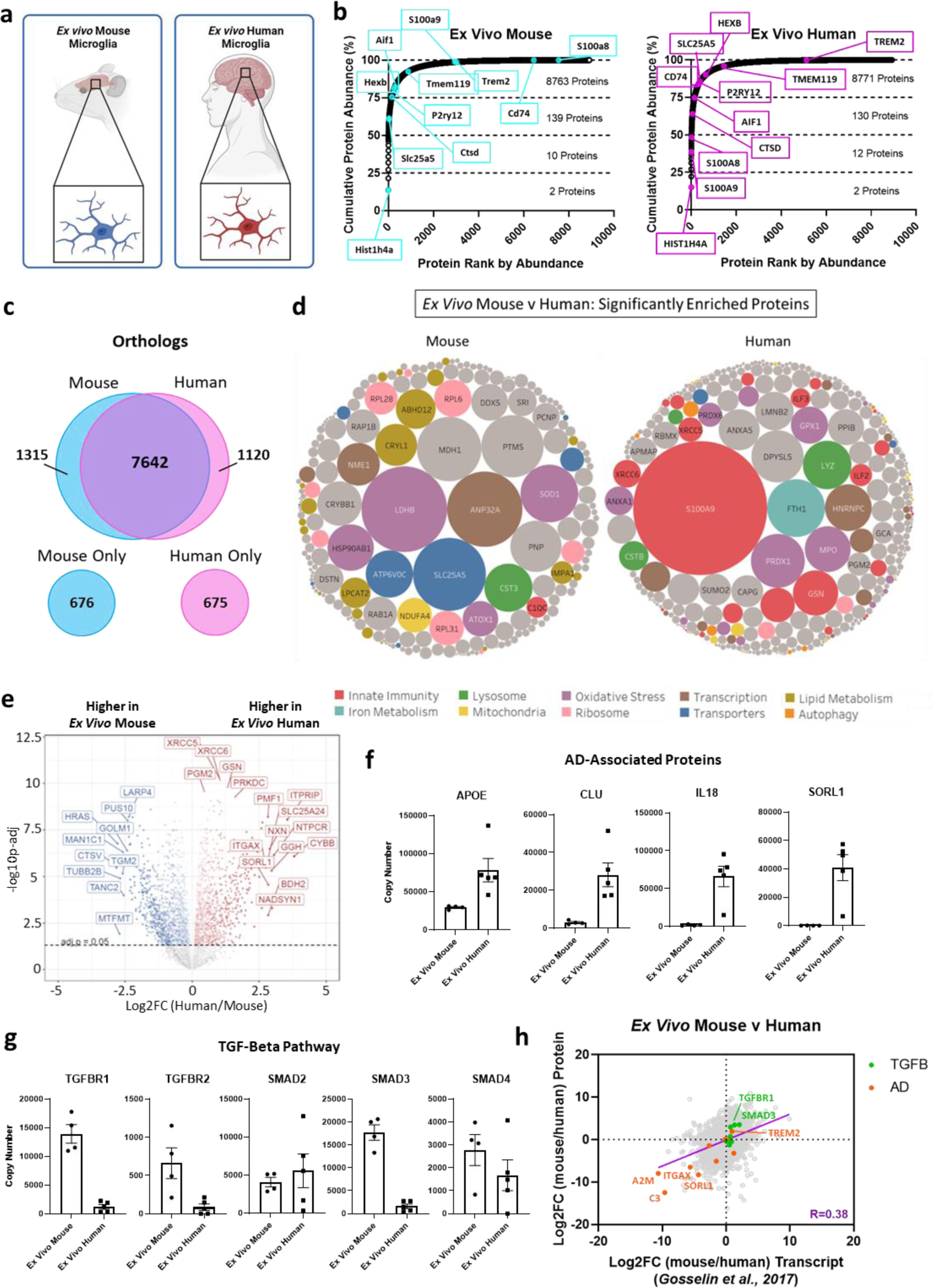
a) Schematic of *ex vivo* mouse and human microglia. **b)** Ranked proteins detected in *ex vivo* mouse and *ex vivo* human microglia based on average copy numbers (x-axis) and cumulative abundance of the total proteome (y-axis). **c)** Venn diagram highlighting shared and unique protein orthologs found in mouse and human proteomes. 676 proteins detected in mouse microglia had no human orthologs and 675 proteins detected in human microglia had no mouse orthologs. **d)** Bubble plots displaying all significantly (p-adj<0.05) differentially expressed proteins of *ex vivo* mouse and human microglia. Proteins are grouped together based on GO Term biological processes. Grey circles represent proteins not associated with any selected GO Terms. Size of circles is proportional to the abundance of each protein (based on copy number). **e)** Volcano plot of differentially expressed proteins (copy number) between *ex vivo* mouse (blue) and human (red) microglia. Proteins that are not significantly differentially expressed (p-adj>0.05) are in grey and below the dotted line. **f)** Total copies per cell of APOE, CLU, IL-18 and SORL1 proteins in *ex vivo* mouse (n=4) and human microglia (n=5) ±SEM. **g)** Total copies per cell of TGF-β proteins TGFBR1, TGFBR2, SMAD2, SMAD3 and SMAD4 in *ex vivo* mouse (n=4) and human microglia (n=5) ±SEM. **h)** Comparison of differentially expressed proteins between *ex vivo* mouse and human microglia at the protein level (y-axis) compared to differentially expressed transcripts between *ex vivo* mouse and human microglia taken from *Gosselin et al., 2017* [28]. Differential expression of both proteins and transcripts is displayed as log2 fold changes. Purple line indicates slope of simple linear regression (0.61). Pearson correlation between protein and transcript log2FC = 0.38 (p<0.0001). Top right quadrant = genes and proteins higher in mouse, bottom left quadrant = genes and proteins higher in human. Proteins of interest are highlighted, associated either with TGF-β pathway (green) or Alzheimer’s Disease (AD; orange).

In both mouse and human, 25% of the proteome composed of two histone proteins (Fig. 3b). with just 12 and 14 proteins making up 50% of the total protein mass for mouse and human microglia, respectively. The top 20 most abundant proteins for human microglia are listed in Supplemental Fig. 3d. Interestingly, the calprotectin proteins S100A9 and S100A8 are within these top 14 proteins in *ex vivo* human microglia (Fig. 3b, Supplemental Fig. 3d) with millions of copies of both proteins detected per cell. Although both proteins are detected in *ex vivo* mouse microglia, they are much less abundant (Fig. 3b). Both subunits, with known functions as both pro-inflammatory and antimicrobial mediators, are known to be expressed by both mouse and human microglia at the transcript level [32] and highly expressed in the brain at the protein level [33], yet their role in microglia is not fully understood. The expression of core microglia markers are similar between mouse and human *ex vivo* microglia.

To compare in more detail protein expression between mouse and human microglia datasets, gene names assigned to proteins were merged on MGI’s human-mouse orthologs on a one-to-one basis with high stringency such that shared mouse and human genes can be mapped together. This identified a total of 11428 proteins across species. Of these proteins, 8762 human proteins had mouse orthologs, and 8957 mouse proteins had human orthologs. Importantly 675 human and 676 mouse proteins had no orthologs at all (Fig. 3c). Gene ontology of species-specific proteins including species-unique orthologs revealed enrichment of complement and antigen presentation in human microglia and translation, DNA replication and cerebral cortex development in mouse (Supplemental Fig. 4a). Unbiased Gene Set Enrichment Analysis (GSEA) of differentially expressed proteins can be found in Supplemental File 1. From a functional point of view, it is important to note that 1120 human proteins with mouse orthologs were not detected in any mouse microglia datasets, and 1315 mouse proteins with human orthologs were not detected in human microglia datasets (Fig. 3c). While 7642 proteins with both mouse and human orthologs are found in both species these differences are important and require consideration when modelling human diseases in mouse models.

To explore this further, we focused on comparing protein expression in *ex vivo* mouse and human microglia proteomes. Of all proteins with shared orthologs, 6921 proteins were expressed by both *ex vivo* mouse and human microglia, making up 66% of all proteins identified in both groups, with one third of all proteins detected only in mouse or human (Supplemental Fig. 4b). AD associated proteins with known mouse and human orthologs [15] were investigated. Out of the 26 AD-associated proteins investigated, 8 were not detected in *ex vivo* mouse nor human microglia samples, and 13 proteins did not significantly differ in their expression (Supplemental Fig. 4c). However, AD-associated proteins ABCA7 and TREM2 were found to be significantly enriched in mouse microglia, and APOE, SCIMP and SORL1 significantly enriched in human microglia.

Differentially expressed proteins further highlighted the increased basally activated state of human microglia compared to mouse by revealing highly abundant enriched human proteins associated with immune responses, including S100A9, GSN, and ILF2 and 3 (Fig. 3d). The full list of proteins can be found in Supplemental File 1. Proteins associated with lipid metabolism were highly abundant enriched proteins in mouse microglia. Many of these proteins show similar differential expression at the transcript level between human and mouse microglia [34].

Further analysis of all differentially expressed proteins (Fig. 3e) revealed significant enrichment of proteins associated with ‘Innate Immune Response’ (Supplemental Fig. 4d) and ‘Alzheimer’s Disease’ in human microglia (Fig. 3f) and ‘Transforming Growth Factor Beta’ (TGF-β) in mouse microglia (Fig. 3g, Supplemental Fig. 4d). COTL1, a microglia-specific marker associated with AD pathology [24], was highly abundant in both mouse and human but not differentially expressed. As TGF-β regulates microglia homeostasis, with loss of signalling associated with primed microglia and activation, [35] enrichment of these networks may highlight the relative reduced exposure of mouse microglia to external stimuli, due to housing conditions and environment, compared to the basally primed state of human microglia. This is further highlighted in enriched protein networks associated with unique and differentially expressed proteins in human microglia heavily associated with immune signalling and inflammation. Indeed, when comparing the Log2 fold changes between *ex vivo* mouse and human microglia at the transcript and protein level, we found these distinct expression profiles are also observed to some extent at the transcript level [28], including the increased expression of TREM2 in mouse microglia (Fig. 3h). Overall correlation between differential transcript and protein was low (Pearson Correlation, R=0.38) but in line with other studies [17–21]. Nevertheless, enrichment of AD- and inflammation-associated proteins in human, and TGF-β proteins in mouse were corroborated by transcriptomics (Fig. 3h). Such differences between mouse and human microglia represent an obvious caveat of animal models particularly in the study of immune responses.

To understand the extent of environment and species influence on these protein expression networks, we investigated the expression of TGF-β and AD-associated proteins in xenograft microglia, a model of human-derived microglia in the mouse brain environment. Interestingly, we detected similar copies of TGFBR1 and 2 proteins in xenograft microglia to mouse, with SMAD2 being most highly expressed in xenograft microglia (Supplemental Fig. 4e). However, we also observed similar levels of APOE, IL-18 and SORL1 in xenograft microglia to *ex vivo* human (Supplemental Fig. 4f), despite the striking differences in environment and exposure. This suggests that expression of these proteins cannot be due to environment alone but may represent species-specific enrichment.

### Mouse microglia undergo rapid morphological and proteomic reshaping in vitro

We next compared the proteomes of mouse microglia directly after sorting from the mouse brain (*ex vivo*) compared to microglia cultured for 7 days (*in vitro*) (Fig. 4a). *In vitro* mouse microglia have triple the protein content of *ex vivo* cells (29pg/cell v 88pg/cell; Fig. 4b). By tracking *in vitro* mouse microglia over the 7-day period, we observed striking morphological changes (Fig. 4c) and a 7-fold increase in cell size from day 0 to day 7 in culture (106µm^2^ v 729µm^2^; Fig. 4d). Increases in protein mass showed a very similar trend to cell size, such that there was a striking increase in both between day 2 and 4, and a steadier increase from day 4 to day 7 (Fig. 4e).

**Figure 4.**
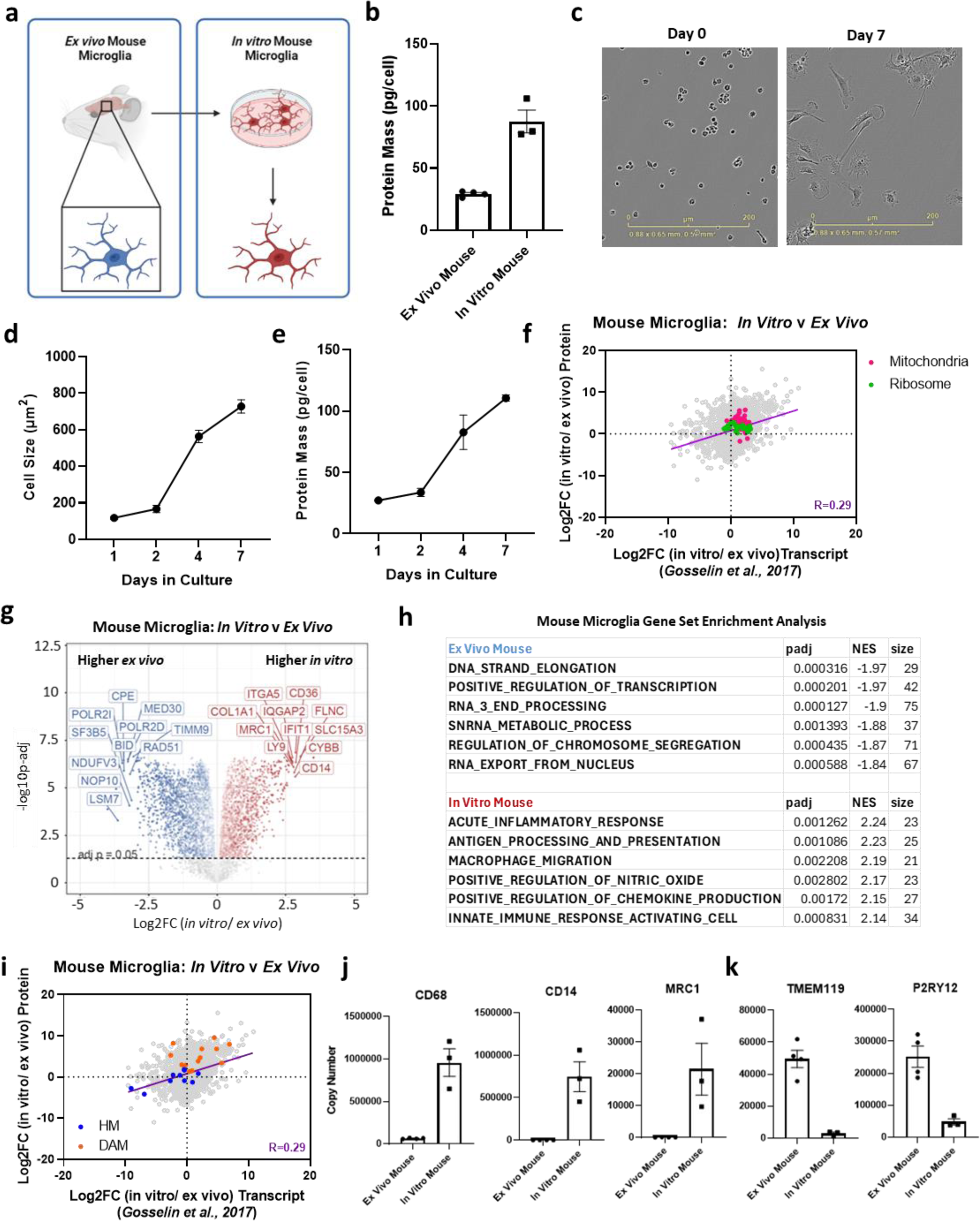
a) Schematic of *ex vivo* and *in vitro* mouse microglia. Samples of *in vitro* mouse microglia were taken after 7 days in culture, created using Biorender. **b)** Protein mass (picograms (pg) per cell) of *ex vivo* and *in vitro* mouse microglia (7 days in culture) ±SEM. N=3. **c)** Representative images of mouse microglia after isolation and seeding (Day 0) and 7 days in culture (Day 7). Scale bar = 200µm. **d)** Median cell size (µm^2^) of mouse microglia after 1, 2, 4 and 7 days *in vitro* ±SEM. N=4. **e)** Protein mass (pg per cell) of mouse microglia after 1, 2, 4 and 7 days *in vitro* ±SEM. N=4. **f)** Comparison of differentially expressed proteins between *in vitro* and *ex vivo* mouse microglia at the protein level (y-axis) compared to differentially expressed transcripts, taken from *Gosselin et al., 2017* [28]. Differential expression of both proteins and transcripts is displayed as log2 fold changes. Purple line indicates slope of simple linear regression (0.46). Pearson correlation between protein and transcript log2FC = 0.29 (p<0.0001). Top right quadrant = genes and proteins higher in *in vitro* mouse microglia, bottom left quadrant = genes and proteins higher in *ex vivo* mouse microglia. Proteins of interest are highlighted, associated either with mitochondria (magenta) or ribosomes (green). **g)** Volcano plot of differentially expressed proteins in *ex vivo* (blue) and *in vitro* (red) mouse microglia. Proteins that are not significantly differentially expressed (p-adj>0.05) are in grey and below the dotted line. **h)** Gene set enrichment analysis (GSEA) of differentially expressed proteins between *ex vivo* and *in vitro* mouse microglia. **i)** Comparison of differentially expressed proteins between *in vitro* and *ex vivo* mouse microglia at the protein level (y-axis) compared to differentially expressed transcripts, taken from *Gosselin et al., 2017* [28]. Differential expression of both proteins and transcripts is displayed as log2 fold changes. Purple line indicates slope of simple linear regression (0.46). Pearson correlation between protein and transcript log2FC = 0.29 (p<0.001). Top right quadrant = genes and proteins higher in *in vitro* mouse microglia, bottom left quadrant = genes and proteins higher in *ex vivo* mouse microglia. Proteins of interest are highlighted, associated either with microglia homeostasis (blue) or DAM (orange). **j)** Total copies per cell of inflammatory proteins CD68, CD14 and MRC1 in *ex vivo* (n=4) and *in vitro* mouse microglia (n=3) ±SEM. **k)** Total copies per cell of homeostatic microglia proteins TMEM119 and P2RY12 in *ex vivo* (n=4) and *in vitro* mouse microglia (n=3) ±SEM.

These independent mouse microglia samples revealed striking reproducibility in proteomic differences when compared with *ex vivo* and *in vitro* mouse datasets. As well as similar increases in protein mass, differential enrichment of inflammatory and microglia identity proteins were also validated in this dataset (Supplemental Fig. 5a).

As increases in nutrient availability can induce mTORC1 signalling [36], we also observed increases in expression of mTORC1 subunits MTOR, MLST8 and RIPTOR, upstream activator RHEB and downstream translation initiator EIF4E (Supplemental Fig. 5b) consistent with the increase in protein mass. This data shows the rapid morphological and proteomic reprogramming of microglia induced by the culture environment over time.

We then compared differentially expressed proteins between *in vitro* and *ex vivo* mouse microglia to transcriptomics data [28] and found enrichment of mitochondrial and ribosomal proteins in *in vitro* mouse microglia were also found at the transcript level (Fig. 4f), although overall correlation between proteins and transcripts was poor (Pearson Correlation, R=0.29). Despite having identical ontogeny, 18% of proteins were not shared between *in vitro* and *ex vivo* microglia (Supplemental Fig. 5c), and ∼32% were differentially expressed (Fig. 4g), revealing significant proteomic reprogramming induced by culture.

To understand if the lower protein mass of *ex vivo* cells is due to fluorescently activated cell sorting (FACS), we performed mass spectrometry analysis of *in vitro* BV2 cells by direct lysis or processing and sorting by FACS (Supplemental Fig. 6a). Sorted cells were either unstained or fluorescently stained to investigate if antibody incubation and therefore prolonged exposure to low (4°C) temperatures also had an effect. FACS did not lead to any differences in the numbers of proteins identified by mass spectrometry (Supplemental Fig. 6b), nor the protein mass (Supplemental Fig. 6c) in line with previous investigations [37]. Furthermore, 99% of all proteins identified were found in all 3 experimental groups (Supplemental Fig. 6d) suggesting that sorting did not impact the detection of proteins. Less than 2% of all proteins were significantly differentially expressed between sorted and unsorted cells, and no significant changes in protein expression were observed between sorted cells that were stained or unstained (Supplemental Fig 6e), suggesting that neither sorting nor prolonged incubation at 4°C for antibody staining significantly impacts protein identification or expression.

GSEA revealed enrichment of protein networks associated with inflammatory responses, antigen presentation and chemokine production in *in vitro* microglia and transcriptional regulation in *ex vivo* microglia (Fig. 4h, full GSEA list in Supplemental File 1). Enrichment of homeostatic proteins were found in *ex vivo* microglia at both the protein and transcript level (Fig. 4i), and although DAM-associated proteins [38] were enriched in *in vitro* microglia proteomes, these were underrepresented at the transcript level (Fig. 4i). Inflammatory mediators CD68, CD14 and MRC1 were significantly enriched in *in vitro* microglia (Fig. 4j) and homeostatic markers TMEM119 and P2RY12 in *ex vivo* microglia (Fig. 4k). Taken together, this data shows the extent of proteomic reshaping that occurs in mouse microglia in culture, with large increases in protein synthesis, energy demand, nutrient sensing and inflammatory protein expression.

### Mouse brain engraftment of human ESC-microglia induces a homeostatic proteomic signature

Having observed the rapid proteomic reprogramming of cells from the *in vivo* to the *in vitro* environment, we investigated the influence of the brain environment on the proteome of previously *in vitro* cultured cells. For this we compared hESC-microglia *in vitro* and after 1-month post-engraftment into the mouse brain. Xenograftment of human derived cells into mice opens the opportunity to model human cellular responses to disease associated stimuli in the tissue environment. This is particularly promising for human brain-resident cells that cannot be routinely obtained. Xenografted hESC-microglia models have been established with full integration of long-lasting hESC-microglia in the brains of immunocompromised Rag^-/-^ mice with humanised CSF to sustain human microglia survival [15, 39]. Importantly, xenografted microglia show a similar transcriptomic profile to *ex vivo* human microglia [15, 39, 40] and respond to disease-associated stimuli such as amyloid beta [15, 41]. We therefore compared the proteomes of hESC-microglia *in vitro* and post-xenograft with *ex vivo* human microglia to determine their similarities at the protein level (Fig. 5a). Indeed, principal component analysis of protein copy number expression in the three human microglia groups prior to differential analysis show close clustering of xenograft and *ex vivo* human microglia compared to *in vitro* hESC-microglia (Fig. 5b), supporting previous evidence of increased similarity of xenografted microglia to *ex vivo* human microglia [15, 40, 42].

**Figure 5.**
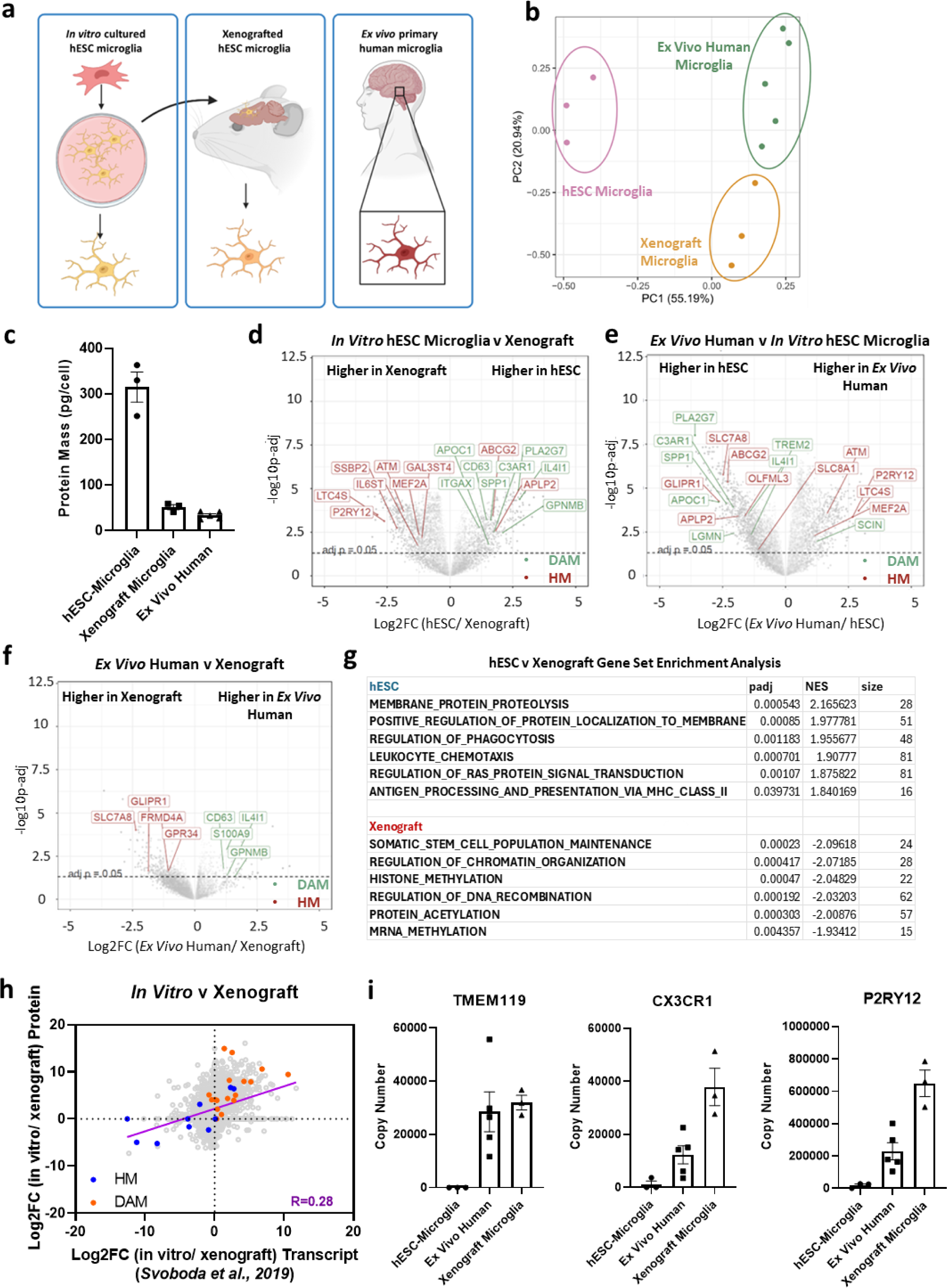
a) Schematic of all human derived microglia groups; *in vitro* cultured hESC-derived microglia, xenografted hESC-microglia and *ex vivo* human microglia. **b)** Principal component analysis (PCA) of *in vitro* hESC-microglia (pink, n=3), xenograft hESC-microglia (orange, n=3) and *ex vivo* human microglia (green, n=5) based on protein copy numbers. Each dot represents a biological replicate for each group. **c)** Protein mass per cell (pg) of *in vitro* hESC-microglia (n=3), xenograft microglia (n=3) and *ex vivo* human microglia (n=5) ±SEM. **d)** Volcano plots displaying the differential expression of proteins between *in vitro* hESC-microglia and xenograft microglia. Proteins highlighted in red are proteins associated with microglia homeostasis (HM) and those in green are associated with disease associated microglia (DAM). Proteins that are not significantly differentially expressed (p-adj>0.05) are in grey and below the dotted line. **e)** Volcano plots displaying the differential expression of proteins between ex vivo human microglia and *in vitro* hESC microglia. Proteins highlighted in red are proteins associated with microglia homeostasis (HM) and those in green are associated with disease associated microglia (DAM). Proteins that are not significantly differentially expressed (p-adj>0.05) are in grey and below the dotted line. **f)** Volcano plots displaying the differential expression of proteins between *ex vivo* human and xenograft microglia. Proteins highlighted in red are proteins associated with microglia homeostasis (HM) and those in green are associated with disease associated microglia (DAM). Proteins that are not significantly differentially expressed (p-adj>0.05) are in grey and below the dotted line. **g)** GSEA of differentially expressed proteins between hESC and xenograft microglia. **h)** Comparison of differentially expressed proteins between *in vitro* hESC-microglia and xenograft microglia at the protein level (y-axis) compared to differentially expressed transcripts comparing hiPSC-derived microglia *in vitro* and after xeno-engraftment from Svodboda et al., 2019 [40]. Purple line indicates slope of simple linear regression (0.47). Pearson correlation between protein and transcript log2FC = 0.28 (p<0.0001). Top right quadrant = genes and proteins higher in *in vitro* stem cell derived microglia, bottom left quadrant = genes and proteins higher in xenograft microglia. Proteins of interest are highlighted, associated either with microglia homeostasis (blue) or DAM (orange). **i)** Total copies per cell of microglia homeostatic proteins TMEM119, P2RY12 and CX3CR1 in *in vitro* hESC-microglia (n=3), *ex vivo* human microglia (n=5) and xenograft microglia (n=3) ±SEM.

Protein content of xenografted hESC-derived microglia drastically decreased from ∼300pg/cell *in vitro* to ∼50pg/cell in the brain environment, much closer in mass to *ex vivo* human microglia than their *in vitro* counterparts (Fig. 5c). Differentially expressed proteins between each group highlighted an enrichment of DAM signature proteins [38] in *in vitro* hESC-microglia (Fig. 5d-f). Compared to *in vitro* hESC-microglia, xenografted microglia displayed an enriched homeostatic signature (Fig. 5d) and importantly, fewer differentially expressed proteins were observed between xenografted microglia and *ex vivo* microglia (Fig. 5f). Despite being the same cell type, 29% of proteins identified were not shared between *in vitro* and xenografted hESC microglia (Supplemental Fig. 7a), demonstrating a large environment-driven proteomic divergence. GSEA between *in vitro* hESC-microglia and xenografted microglia reveal enrichment of RAS signalling, antigen presentation and phagocytosis in hESC-microglia, and histone methylation, protein acetylation and transcription in xenograft microglia (Fig. 5g, full GSEA list in Supplemental File 1). Importantly, the brain environment induced expression of markers associated with microglia identity and homeostasis in hESC-microglia; GSEA also highlighted enrichment of DAM proteins in the *in vitro* hESC-microglia, and homeostatic proteins in xenograft microglia (Supplemental Fig. 7b, c). We compared our data to published transcriptomics data comparing hiPSC-microglia *in vitro* and post-engraftment into the mouse brain [40]. Although enrichment of homeostatic markers was observed at both protein and transcript levels in xenografted microglia, DAM-enrichment observed in *in vitro* human stem cell derived microglia was much more prominent again at the protein level (Fig. 5h). Once again, overall correlations between differentially expressed protein and transcripts were poor (R=0.28).

Finally, we analysed the expression of homeostatic microglia markers TMEM119, P2RY12 and CX3CR1 (Fig. 5i) and observed that although these markers were not detected or very lowly abundant in *in vitro* hESC-microglia, engraftment into the mouse brain led to increase expression of all three proteins to comparable levels to *ex vivo* human microglia. This data further identifies the influence of environment on proteomic programming of microglia cells, and the particular advantages of xenograft models in inducing a more homeostatic and ‘human-like’ proteome in hESC-microglia.

## Discussion

To understand the role of microglia in health and disease, it is imperative that we understand the advantages and limitations of commonly used model systems. This deep proteomic analysis gives detailed insight into the differential phenotypes of these cells including quantifiable expression of proteins relating to metabolism, environmental sensors, and inflammatory profile. Our dataset showcases fundamental proteomic differences of mouse and human, and *ex vivo* and *in vitro* microglia. Comparisons to published transcriptomic datasets highlight the disparity between transcript and protein expression, in particular, the lack of DAM-associated marker enrichment induced by *in vitro* culturing in both mouse and human microglia, by transcriptomics. These comparisons highlight the importance of defining phenotypes and functions via protein expression. Our data enriches our understanding of human and mouse microglia biology and the plasticity of proteomic reprogramming that microglia undertake in response to environmental changes, as well as providing a detailed proteomic map of microglia as an important resource for the field.

Our database includes the BV2 cell line proteome, a commonly used *in vitro* model to analyse microglia-like responses. The BV2 cell line is derived from primary mouse (C57/BL6) microglia that have been immortalised using v-raf/v-myc carrying J2 retroviruses. Their rapid proliferation and immortalisation provide many advantages over primary microglia culture, however their suitability as a model of microglia responses has been debated. Our data has highlighted the BV2 cell line as proteomically distinct from all other microglia groups, and therefore we have omitted this data from our main comparisons. We do believe however that it is essential that we provide a detailed proteome for the field to interrogate and understand their limitations, available in Supplemental File 1.

Limitations of human microglia studies warrants the need to use appropriate animal models to investigate the role of microglia in health and disease, yet disparities between mouse and human microglia transcriptomes, particularly in the context of neurodegeneration [16, 20] and ageing [43] have been observed. Enrichment of DAM proteins also differs in microglia from AD patient brains and mouse models [38, 44], although how unstimulated microglia proteomes compare is less understood. Our data highlights interesting similarities and differences between human and mouse microglia. Although glycolysis represented a much lower proportion of the proteome than ribosomal or mitochondrial proteins for both mouse and human microglia, there was a 4-fold higher proportion in *ex vivo* mouse microglia compared to human. This was surprising considering human microglia displayed enrichment of inflammatory proteins which is also associated with metabolic shifts from oxidative phosphorylation to glycolysis [45, 46]. Inhibition of glycolysis with 2-doxyglucose leads to cell death in mouse macrophages but not human [46], therefore may represent an increased basal glycolytic demand of mouse microglia.

Human microglia showed significant enrichment of proteins associated with inflammation compared to mouse microglia. Contributing factors associated with these distinct protein expression patterns include age, environmental exposures, and inherent species differences. Primary human microglia are often obtained from surgically resected tissue derived from either epilepsy or brain tumour patients using established pathological characterisation of non-pathological tissue and isolation methods [13, 15, 27-29]. Although adjacent to pathology, microglia isolated in this way are from non-pathological areas, with previous characterisation identifying them to have a highly homeostatic gene signature [13, 15]. Indeed, expression of homeostatic marker P2RY12 is comparable between adult human microglia isolated from non-pathological areas of tissue and foetal human microglia at the transcript level [29]. Furthermore, transcriptomic analysis of microglia from the centre of epilepsy pathology [47] compared to microglia isolated from non-pathological adjacent tissue [13] showed distinct gene expression patterns in epilepsy tissue-resident microglia consistent with increased inflammation and antigen presentation. Importantly, high IL-1β expression was observed in epilepsy microglia compared to non-diseased controls (33.5% of cells vs 6.9%), consistent with previous findings that demonstrated increased IL-1β expression in microglia from mouse models of pilocarpine-induced epilepsy [48]. We do not detect IL-1β in any of our *ex vivo* human microglia samples (Supplemental File 1), supporting the argument that microglia from non-pathological tissue adjacent to pathology do not have an epilepsy-associated gene or protein signature. We are therefore confident that proteomic differences between *ex vivo* mouse and human are not due to an epilepsy phenotype in our human microglia samples.

Fundamental differences in expression of key AD-related proteins was also shown, with a surprising finding of TREM2 enrichment in *ex vivo* mouse microglia. TREM2 is an important example highlighting the limitations of mouse models in the cellular and molecular study of neurodegenerative disease. TREM2 loss of function mutations are associated with Nasu-Hakola disease in humans, characterised by early onset dementia and bone cysts, yet this does not manifest in mice [49]. TGF-β pathway enrichment in mouse microglia was evident, along with a notable lack of inflammatory proteins detected, suggestive of a more naïve and homeostatic microglia state [35, 50] compared to human microglia. However, mouse microglia have been shown to express greater levels of Tgfb1 transcripts compared to both adult and foetal human microglia [28, 29], suggesting that TGF-β expression may be species-specific.

Investigating both AD-associated and TGF-β pathway proteins in xenograft microglia gave us some insight into species and environmental driven expression of proteins. We found that these cells had high expression of both TGF-β signalling and AD-associated proteins. This suggests that high human microglia expression of proteins associated with AD cannot be due to age and exposure alone, as xenograft cells are derived from sterile *in vitro* culture and xenografted into young adult mice in a clean housing environment. H9 stem cells used for these studies have an APOE3/4 genotype and a low polygenic risk score for AD (0.4) [51], therefore we would not expect an AD phenotype or AD-associated protein signature in these cells. Conversely, expression of TGF-β proteins in xenograft microglia may highlight the influence of the environment on proteomic programming.

Altogether, this data highlights fundamental differences in protein expression, some of which are known to differ in their downstream signalling, all of which may contribute to the divergent microglia inflammatory phenotypes and responses to stimuli in experimental models.

One of the most striking differences between *in vitro* and *ex vivo* microglia is their protein mass and regulation of translation, as shown by EIF4A1:PDCD4 ratio. This does not occur in all immune cells, as naïve CD4 and CD8 T-cells in culture maintain tight regulation of translation [30], with reduction in PDCD4-regulation of EIF4A1 only occurring after stimulation. *In vitro* cultured microglia display an activated phenotype without stimulation, as shown by downregulation of homeostatic markers TMEM119, CX3CR1 and P2RY12, increase in metabolic demand and inflammatory protein production. Indeed, microglia responses to stimuli are known to differ *in vitro* and *in vivo* [25, 26] with exaggerated responses observed in culture [26]. Heightened inflammatory responses are also observed in aged microglia combined with a primed inflammatory phenotype [52, 53], also observed in our *in vitro* microglia datasets. Furthermore, aged mouse microglia share a transcriptomic profile similar to DAM [53] again consistent with our *in vitro* findings of a pronounced DAM proteomic profile particularly evident in hESC-microglia which significantly decreases after xenograftment. It is therefore perhaps unsurprising that protein modules enriched in the *in vitro* microglia share commonalities with aged microglia, such as increased mTOR signalling components.

Nutrient abundance in culture media can promote mTOR activity [54], initiating the production of proteins, lipids and nucleotides whilst inhibiting autophagy and other catabolic processes. Although a regulator of cap-dependant translation, mTOR appears to show bias in promoting the translation of inflammatory proteins [52], thought to be responsible for the increase in basal inflammatory protein expression of cells *in vitro*. This data shows the extent of the culture environment’s influence on microglia phenotypes, and the important considerations needed when assessing phenotypic changes of microglia *in vitro*.

An interesting feature of our microglia datasets is that plasticity of proteomic reprogramming of cells changing their environment, and crucially, the rescue of the *in vitro* microglia phenotype once engrafted into the brain. This has recently been observed in alveolar macrophages, where even long term culture-induced epigenetic changes associated with increased glycolytic activity, cell migration and adhesion, were completely reversed upon engraftment back into the lung [55]. Xenografted human stem cell derived microglia share large transcriptional similarities to *ex vivo* human microglia [15, 19] which is now corroborated at the protein level. With divergent responses to inflammatory and disease-associated stimuli between mouse and human microglia [13, 20], xenografts represent an exciting avenue of research to understand human microglia responses in tractable disease models amenable to genetic and pharmacological manipulation.

An important factor in microglia homeostasis is maintaining neuronal and glial cellular contact. *In vitro* cultured iPSC-derived microglia maintain expression of homeostatic markers and decrease basal inflammatory protein expression when co-cultured with neurons and astrocytes [56, 57]. It would therefore be interesting to analyse the proteomic signatures of microglia in culture conditions such as co-cultures, organoids, or slice cultures where cellular contacts are maintained, to understand if this alone can rescue the inflammatory phenotype of monocultured microglia.

Overall this data has generated in-depth proteomic maps of human and mouse microglia from *in vitro* and *ex vivo* sources, and provided greater insight into the proteomic reprogramming that occurs *in vitro*. We provide a detailed resource of microglia proteomes for the field to accelerate our understanding of microglia biology.

## Methods

### Human Samples and Isolation of Primary Human Microglia by FACS

Human primary microglia were isolated from brain tissue samples resected from the temporal cortex during neurosurgery (n=5 female, ages 6-61). Patient information is detailed in Supplemental Fig. 2c and Supplemental File 1. All samples represented lateral temporal neocortex and were obtained from patients who underwent amygdalo-hippocampectomy for mesial temporal lobe seizures. The mesial temporal specimens were sent to pathology and thus not available for study purposes. Samples were collected at the time of surgery and immediately transferred to the lab for tissue processing, with post sampling intervals of 5-10 min. All procedures were conducted to protocols approved by the local Ethical Committee (protocol number S61186). Brain biopsies were placed in PBS 2%, FCS, 2mM EDTA (FACS buffer) with Golgi Plug (Brefeldin A, 420601, BioLegend), mechanically triturated and enzymatically dissociated using the Neural Tissue Dissociation Kit (P) (Miltenyi) following manufacturer specifications. Then, samples were passed through a cell strainer of 70μm mesh (70µm strainers, Grenier BioOne) with FACS buffer, and centrifuged at 300g for 15 min at 4°C. Next, cells were resuspended in 30% Percoll (100% Percoll Plus, 17-5445-02, GE Healthcare) and centrifuged at 300g for 15 min at 4°C. The supernatant and myelin layers were discarded, and the cell pellet was treated to remove red cells (Red blood cell lysis buffer, 420301, Biolegend). After a wash, antibody labelling was performed for 30 min at 4°C, using the anti-CD11b (1:50, 130-113-806, Miltenyi Biotec) and anti-CD45 (1:50, 555485, BD Bioscience), adding e780 (1:2000, 65-0865-14, ThermoFisher) as a cell viability marker. Samples were run on a MACSQuant Tyto cell sorter and data were analysed using FCS express software. Human cells were sorted according to the expression of CD11b, CD45. After isolation, cells were washed two consecutive times with ice cold HBSS, snap frozen, and stored at −80°C.

### Mice

Male C57BL/6J mice, aged 3-4 months were used to generate *in vitro* (n=3) and *ex vivo* (n=4) mouse microglia proteomes. For xenotransplant experiments, both male and female Apphu Rag2−/− IL2rγ−/−hCSF1KI were used (n=3, 2 x female, 1 x male). In brief, Rag2−/− IL2rγ−/−hCSF1KI mice, originally purchased from Jacksons Labs (strain 017708), were crossed with Appem1Bdes mice (MGI ID: 6512851) (abbreviated here Apphu) [58]. All animals were bred and housed in groups of 2-3, under a 12h light/dark cycle at 21°C, with food and water ad libitum in local facilities at KU Leuven. All experiments were conducted according to protocols approved by the local Ethical Committee of Laboratory Animals of the KU Leuven (government license LA1210579, ECD project numbers: P177/2020 and P177/2017) following local and EU guidelines.

### Isolation of *Ex vivo* Mouse Microglia by FACS

Mouse primary microglia isolation was performed as previously described [15]. Briefly, mice were terminally anesthetized with an overdose of sodium pentobarbital and transcardially perfused with heparinized PBS. Brains were collected in ice-cold FACS buffer (PBS with 2% FCS and 2mM EDTA). Tissue was mechanically and enzymatically dissociated using the Neural Tissue Dissociation Kit (P) (130-092-628, Mylteni) following manufacturer’s instructions. Next, cell suspension was filtered on a 70-μm cell strainer (542070, Greiner) with FACS buffer and centrifuged at 300g for 15 min at 4°C. Cell pellet was further resuspended in 30% Percoll (GE Healthcare) and spun at 300g for 15 min at 4°C. Myelin cleaning was performed by removing myelin layers formed on top of the supernatant with Pasteur pipettes. Then, cell pellet was incubated with fluorophore-conjugated antibodies for 30 minutes at 4°C using anti-mouse CD11b-PE (1: 50, REA592, Miltenyi), anti-mouse CD45-BV421 (1:500, 563890, BD Biosciences) and the cell viability marker e780 (1:2000, 65-2860-40 eBioscience). Microglia were sorted on a Miltenyi MACSQuant Tyto flow cytometer. ∼1.5 million sorted microglia (CD11b+/CD45m/e780-) from every two mice were pooled, washed with HBSS and snap frozen for the downstream experiments.

### Isolation and Culture of *In vitro* Mouse Microglia

For cell culture, microglia were isolated using MACS Neural Dissociation Kit (Miltenyi) according to manufacturer’s instructions. After perfusion and brain tissue digestion as detailed above, cell pellet was resuspended in MACS buffer (PBS, 0.5% low endotoxin BSA (Merck), 2 mM EDTA, 90μL/brain). Microglia were isolated using an Anti-P2Y12 MoJoSort kit (BioLegend) according to manufacturer’s instructions. Briefly, anti-P2Y12 antibody was added (10μL/brain) and the suspension incubated for 15 min at 4 °C. Cells were washed with MACS buffer, resuspended in 90μL/brain MACS buffer, then streptavidin beads (10μL/brain) were added and incubated for a further 15 mins. Cells were washed again as above then run through pre-rinsed (MACS buffer) LS columns attached to a magnet (Miltenyi). After washing with 12mL MACS buffer, columns were removed from the magnet and cells retained (microglia) were flushed in 5mL MACS buffer. Microglia were resuspended in Dulbecco’s Modified Eagle’s Medium/Nutrient Mixture F-12 (DMEM/F-12, 10565018, ThermoFisher) supplemented with 100 U/mL penicillin and 100μg/mL streptomycin (PenStrep, Merck), 10% heat-inactivated fetal bovine serum (FBS, ThermoFisher), 500ng/mL rhTGFβ-1 (Miltenyi), 10ng/μL mCSF1 (R&D Systems). Microglia were counted using a haemocytometer and plated out onto 96-well plates (Corning). Cells were cultured for 7 days with a half media change on day 3. Cells were collected by direct lysis on plate in 5% SDS, pipetted into 1.5ml eppendorfs, snap frozen and stored at −80°C.

### Culture of BV2 cells

BV2 cells were cultured in RPMI 1640 (Gibco) with 10% foetal bovine serum (FBS) at 37°C with 5% CO_2_, with passaging every 2-3 days with trypsin/EDTA (Gibco). 3 biological replicates were generated for proteomic analysis, each with 1×10^6^ cells per sample. Cells were collected by direct lysis on plate in 5% SDS, pipetted into 1.5ml eppendorfs, snap frozen and stored at −80°C.

### Generation of Human-derived Microglia from ESCs

Human microglia/macrophage precursors were generated from human embryonic stem cells (H9) using the MIGRATE protocol [42]. On days 25 and 32, precursor cells and embryoid bodies (EBs) were harvested and passed through a sterile reversible cell strainer (37μm). Only precursor cells will pass through the strainer and EBs will be retained and re-plated for later collections as detailed [42]. Precursor cells were centrifuged for 5 min at 300 g. Pelleted cells were further resuspended and plated in glass coverslips in a seeding concentration of 10^6 cells/well in microglia differentiation media (TIC) as previously described [15] and based on previous protocols (Abud et al., 2017; Bohlen et al., 2017). TIC media was composed of phenol-red free DMEM/F12, N-acetylcysteine (5μg/ml), insulin (1:2,000), Apo-Transferrin (100μg/ml), sodium selenite (100ng/ml), cholesterol (1.5μg/ml) and heparan sulfate (1μg/ml) supplemented with interleukin-34 (50ng/ml), macrophage colony stimulating factor (M-CSF) (50ng/ml), CX3CL1 (10ng/ml) and transforming growth factor-β (TGF-β; 25ng/ml). Microglia were differentiated in TIC media for 7 days and fully refreshed 3 days after plating. To harvest human-derived microglia for proteomics, on day 7 of differentiation, TIC media was removed and cells were washed twice with DPBS. Next, cells were directly lysed on the plate with 5% SDS in DPBS, scrapped off, transferred to a 1.5ml Eppendorf, snap frozen and stored at −80°C.

### Transplantation of Human Microglia into the Mouse Brain

Generation and grafting of human-derived microglia was performed as previously described [15, 42]. In brief, day 18 microglia/macrophage progenitors were collected and grafted into ApphuRag2−/−IL2rγ−/−hCSF1KI mice brains at a concentration of 250,000 cells/μl at postnatal day 4 (P4). After injections, mice were left recovering in a heating pad and transferred back to their cages. After 10 weeks, mice were sacrificed, perfused and brain tissue processed for FACS as described above and previously [15]. Samples were stained with the following antibodies: CD11b-PE (1:50, REA592, Miltenyi), anti-human CD45 (1:50, 555485, BD Biosciences), anti-mouse CD45-BV421 (1:500, 563890, BD Biosciences) and cell viability marker e780 (1:2000, 65-2860-40 eBioscience). Microglia were sorted on a Miltenyi MACSQuant Tyto flow cytometer. Between 8.5×10^5 and 1.3×10^6 microglia were sorted per sample (CD11b+/hCD45+/mCD45-/e780-), washed with HBSS and snap frozen for downstream proteomic processing.

### Proteomics Sample Preparation

All samples were lysed in 400µl lysis buffer (5% SDS, 10mM TCEP, 50mM TEAB) and shaken at RT for 5 minutes at 1000rpm, followed by boiling at 95°C for 5 minutes at 500rpm. Samples were then shaken again at RT for 5 minutes at 1000rpm before being sonicated for 15 cycles of 30 seconds on/ 30 seconds off with a BioRuptor (Diagenode). Benzonase was added to each sample and incubated at 37°C for 15 minutes to digest DNA. Samples were then alkylated with 20mM iodoacetamide for 1 hour at 22°C. Protein concentration was determined using EZQ protein quantitation kit (Invitrogen) as per manufacturer instructions. Protein isolation and clean up was performed using S-TRAP™ (Protifi) columns before digestion with trypsin at 1:20 ratio (enzyme:protein) for 2 hours at 47°C. Digested peptides were eluted from S-TRAP™ columns using 50mM ammonium bicarbonate, followed by 0.2% aqueous formic acid and 50% aqueous acetonitrile containing 0.2% formic acid. Eluted peptides were dried down overnight.

### Peptide Fractionation

Dried peptides were rehydrated in 210µl 5% formic acid, shaking at 1000rpm for 1 hour at 30°C. Samples were loaded into the high-performance liquid chromatographer (HPLC) with the following buffers: buffer A (10mM ammonium formate with 2% acetonitrile in Milli-Q water (v/v)) and buffer B (10mM ammonium formate with 80% acetonitrile in Milli-Q water (v/v)). Samples were fractionated using high pH reverse phase liquid chromatography using a Dionex Ultimate3000 system. Peptides were loaded onto a 2.1mm x 150mm XBridge Peptide BEH C18 column with 3.5μm particles (Waters) and were separated using a 25 min multistep gradient of solvents A (10mM formate at pH 9 in 2% acetonitrile) and B (10 mM ammonium formate pH 9 in 80% acetonitrile), at a flow rate of 0.3 mL/min. Samples were fractionated into 16 samples and collected into 96 well plates, with blank samples run between samples. All samples were then dried down overnight before resuspension in 5% formic acid.

### Mass Spectrometry

For each sample, 2µg peptide was analysed. Peptides were injected onto a nanoscale C18 reverse-phase chromatography system (UltiMate 3000 RSLC nano, Thermo Scientific) and electrosprayed into an Q Exactive™ Plus Hybrid Quadrupole-Orbitrap™ Mass Spectrometer (Thermo Fisher). For liquid chromatography the following buffers were used: buffer A (0.1% formic acid in Milli-Q water (v/v)) and buffer B (80% acetonitrile and 0.1% formic acid in Milli-Q water (v/v). Samples were loaded at 10 μL/min onto a trap column (100μm × 2cm, PepMap nanoViper C18 column, 5μm, 100 Å, Thermo Scientific) equilibrated in 0.1% trifluoroacetic acid (TFA). The trap column was washed for 3 min at the same flow rate with 0.1% TFA then switched in-line with a Thermo Scientific, resolving C18 column (75μm × 50cm, PepMap RSLC C18 column, 2μm, 100 Å). Peptides were eluted from the column at a constant flow rate of 300nl/min with a linear gradient from 3% buffer B to 6% buffer B in 5 min, then from 6% buffer B to 35% buffer B in 115 min, and finally to 80% buffer B within 7 min. The column was then washed with 80% buffer B for 4 min and re-equilibrated in 3% buffer B for 15 min. Two blanks were run between each sample to reduce carry-over. The column was kept at a constant temperature of 50°C.

Each sample was subject to rigorous quality control before, during and after runs using a HeLa peptide standard with a minimum of 2000 proteins detected. If less than 2000 proteins were detected during a QC run, the machine was cleaned and re-checked before proceeding.

The data was acquired using an easy spray source operated in positive mode with spray voltage at 2.445kV, and the ion transfer tube temperature at 250°C. The MS was operated in DDA mode. A scan cycle comprised a full MS scan (m/z range from 350-1650), with RF lens at 40%, AGC target set to custom, normalised AGC target at 300, maximum injection time mode set to custom, maximum injection time at 20 ms and source fragmentation disabled. MS survey scan was followed by MS/MS DIA scan events using the following parameters: multiplex ions set to false, collision energy mode set to stepped, collision energy type set to normalized, HCD collision energies set to 25.5, 27 and 30, orbitrap resolution 30000, first mass 200, RF lens 40, AGC target set to custom, normalized AGC target 3000, maximum injection time 55ms.

### Proteomics Data Handling, Processing and Analysis

The data were processed, searched and quantified with the MaxQuant software package (Version 1.6.10.43). Raw mass spec data files for human microglia were searched against a human SwissProt database with isoforms (July 2020) while mouse microglia samples were searched against a SwissProt database with isoforms combined with mouse TrEMBL entries with protein level evidence available and a manually annotated homologue within the human SwissProt database (June 2020). The false discovery rate was set to 1% for positive identification at the protein and peptide-to-spectrum match level. Protein N-terminal acetylation and methionine oxidation were set as variable modifications and carbamidomethylation of cysteine residues was selected as a fixed modification. Match between runs was disabled. The dataset was filtered to remove proteins categorized as ‘contaminants’, ‘reverse’ and ‘only identified by site’. To normalise the data across all runs, protein copy numbers were calculated from protein raw intensities using the Histone Ruler method, as described previously [59] using Perseus software (MaxQuant) [60]. Microglia samples were collected, processed and analysed by mass spectrometry over a 2-year period, which increases the potential for batch effects. To study this we used data collected from QC runs composed of a commercial HeLa standard. LFQ intensity data from HeLa runs was used to calculate potential batch effects using the coefficient of variation (CV) which were stable throughout batches, with the exception of the *ex vivo* mouse batch due to three outliers. However, using the Levene test for homogeneity of variation, we concluded that batch effects across HeLa runs were non-significant (P=0.9061), and therefore did not perform any batch corrections on microglia proteome data.

### Statistical Analysis

P values were calculated via a two-tailed, unequal variance t-test on log-normalized data and adjusted using the Benjamini-Hochberg procedure. Proteins with adjusted P values of less than 0.05 were considered significant. The mass of individual proteins was estimated using the following formula: CN×MW/N_A_ =protein mass (g cell^−1^), where CN is the protein copy number, MW is the protein molecular weight (in Da) and N_A_ is Avogadro’s Constant. Heat maps were generated using the Morpheus tool from the Broad Institute (https://software.broadinstitute.org/morpheus). PCA plots and Venn diagrams were generated in R studio. Volcano plots in Supplementary Figure 3a and b were generated using Graphpad Prism. All other volcano plots were generated in R studio using the Limma-Voom package. Weighted Gene Correlation Network Analysis (WGCNA) was performed in R on the copy numbers data in an unsigned network fashion. Resulting modules were annotated via DAVID functional analysis, labelling only modules with significant (adjusted p-value < 0.05) and positive enrichment of at least one GO term.

## Supporting information

Supplemental File 1 Updated

## Data Availability

Raw mass spectrometry files and results files are available through ProteomeXchange using the following login details: username, password 8qstGL56 for the mouse data, username, password 0luxtTJq for the human data and username, password: K8BEE51Q for HeLa standards data.

## Acknowledgements

This work was supported by the UK-DRI grant contributed by MRC (UK). Further funding from Fonds voor Wetenschappelijk Onderzoek (FWO, Belgium), ‘‘Methusalem’’ and ‘‘Opening the Future’’ grants from KU Leuven (Belgium), Stichting Alzheimer Onderzoek (Belgium), the Alzheimer Association (USA), the European Research Council ERC-CELLPHASE_AD834682 (EU), Geneeskundige Stichting Koningin Elisabeth (Belgium), Bax-Vanluffelen (Belgium) and Mission Lucidity (Leuven, Belgium). Schematics in all figures were generated with BioRender.com.

## Author Contributions

A.F.L, B.D.S and A.J.M.H conceived and designed the study. A.F.L carried out the experiments, analysed and interpreted the data and wrote the manuscript. A.M.M generated *in vitro* hESC-derived microglia for proteomics, carried out xenograft experiments and generated xenograft microglia for proteomics. E.D assisted in data analysis, generation of figures and development of online database. M.J.D isolated and cultured *in vitro* mouse microglia for fractionated proteomes. P.H isolated *ex vivo* mouse microglia for proteomics. R.M, A.S and K.C isolated *ex vivo* human microglia for proteomics. A.J.B assisted in data analysis and manuscript preparation. I.G assisted with generating xenograft and *in vitro* hESC-derived microglia samples for proteomics. T.T carried out amygdalo-hippocampectomy surgeries which generated *ex vivo* human microglia samples. M.F assisted in data analysis, data interpretation and development of online database. B.D.S co-supervised the project, co-designed the study and guided experiment design, data interpretation and manuscript preparation. A.J.M.H co-supervised the project, co-designed the study and guided experiment design, data interpretation and manuscript preparation. B.D.S raised the funding. All authors read and approved the manuscript.

---

**Supplemental Figure 1.**
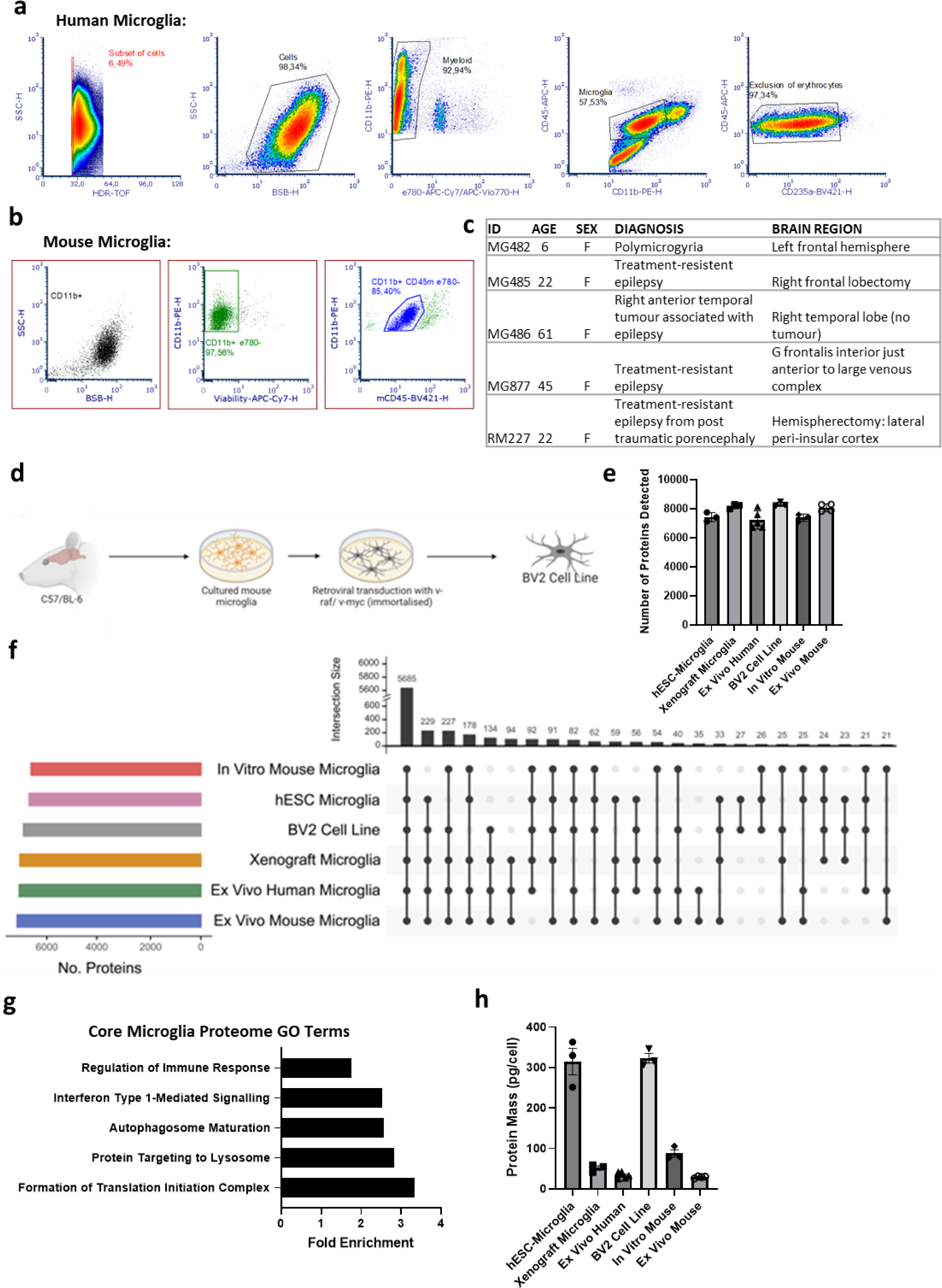
**a)** Gating strategy for the isolation of *ex vivo* human microglia. **b)** Gating strategy for the isolation of *ex vivo* mouse microglia. **c)** Donor information from *ex vivo* human microglia samples. **d)** Schematic of BV2 cell origins. **e)** Total number of proteins detected in all microglia groups and including BV2 cells (n=3) ±SEM. **f)** Upset plot showing the numbers of proteins intersecting between all six microglia groups and highlighting the proteomic convergence for each. **e)** GO terms derived from protein expression across all microglia groups. **g**) GO terms derived from core microglia protein expression across all samples. **h)** Total copies of all histone proteins detected in all six microglia groups ±SEM.

**Supplemental Figure 2.**
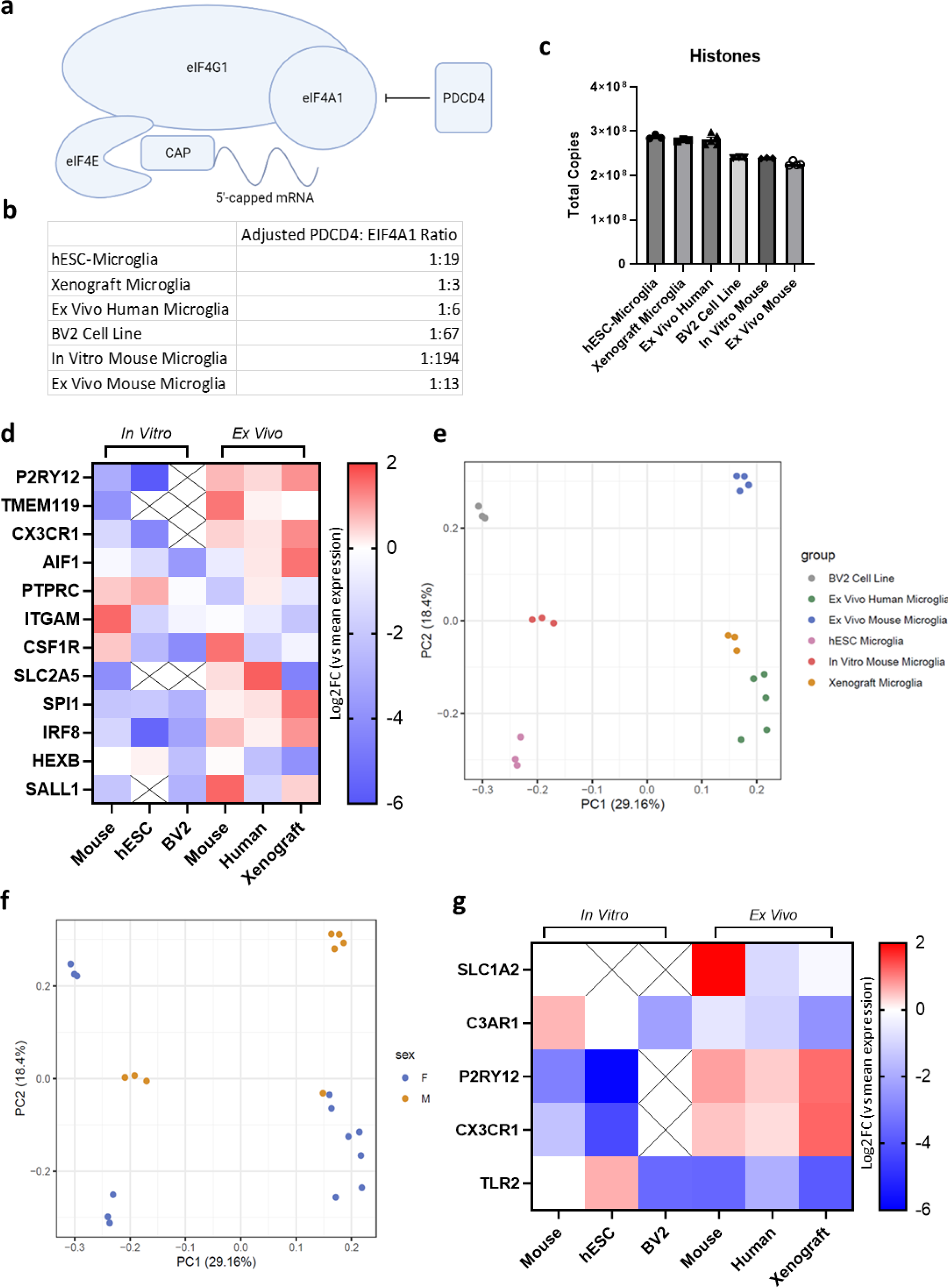
**a)** Schematic of the eukaryotic initiation factor subunit assembly for protein translation. **b)** Adjusted ratios of numbers of PDCD4 proteins for each EIF4A1 protein, an indicator of translation regulation. The lower the ratio, the greater the control of translation. **c)** Total protein mass per cell of all microglia groups and including BV2 cells (n=3) ±SEM. **d)** Relative expression of microglia markers across all samples normalised to the mean expression across all samples, plotted as log2 fold change over the mean. Red = higher than the mean expression, blue = lower than the mean expression. Crossed out boxes indicate no protein detected. **e)** PCA of all microglia groups and the BV2 cell line. Each dot represents a replicate from BV2 cell line (n=3, grey), *ex vivo* human microglia (n=5, green), *ex vivo* mouse microglia (n=4, blue), *in vitro* hESC-derived microglia (n=3, pink), *in vitro* mouse microglia (n=3, red) and xenograft microglia (n=3, yellow). **f)** PCA of all samples identified by their sex, highlighted as either female (blue) or male (orange). **g)** Relative expression of proteins associated with microglia the sensome normalised to the mean expression across all samples, plotted as log2 fold change over the mean. Red = higher than the mean expression, blue = lower than the mean expression. Crossed out boxes indicate no protein detected.

**Supplemental Figure 3.**
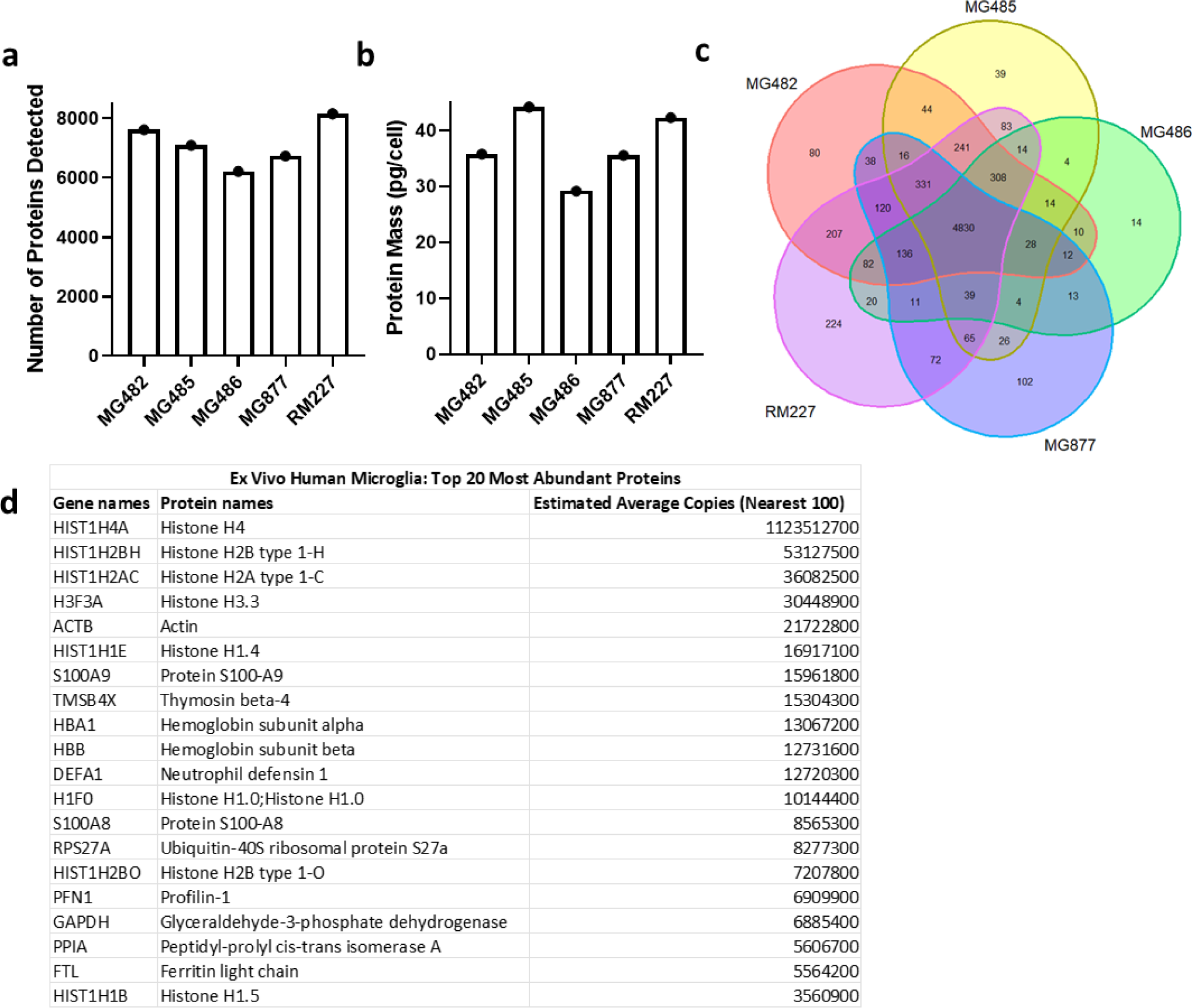
**a)** Total number of proteins detected for each *ex vivo* human microglia sample. **b)** Protein mass (pg per cell) for each *ex vivo* human microglia sample. **c)** Venn diagram of each protein identified in each *ex vivo* human microglia sample, highlighting the similarities and variation between samples. **d)** Table of the top 20 most abundant proteins found in *ex vivo* human microglia calculated using the average copy number per cell (rounded to the nearest 100) of each protein across all samples and ranking by most abundant.

**Supplemental Figure 4.**
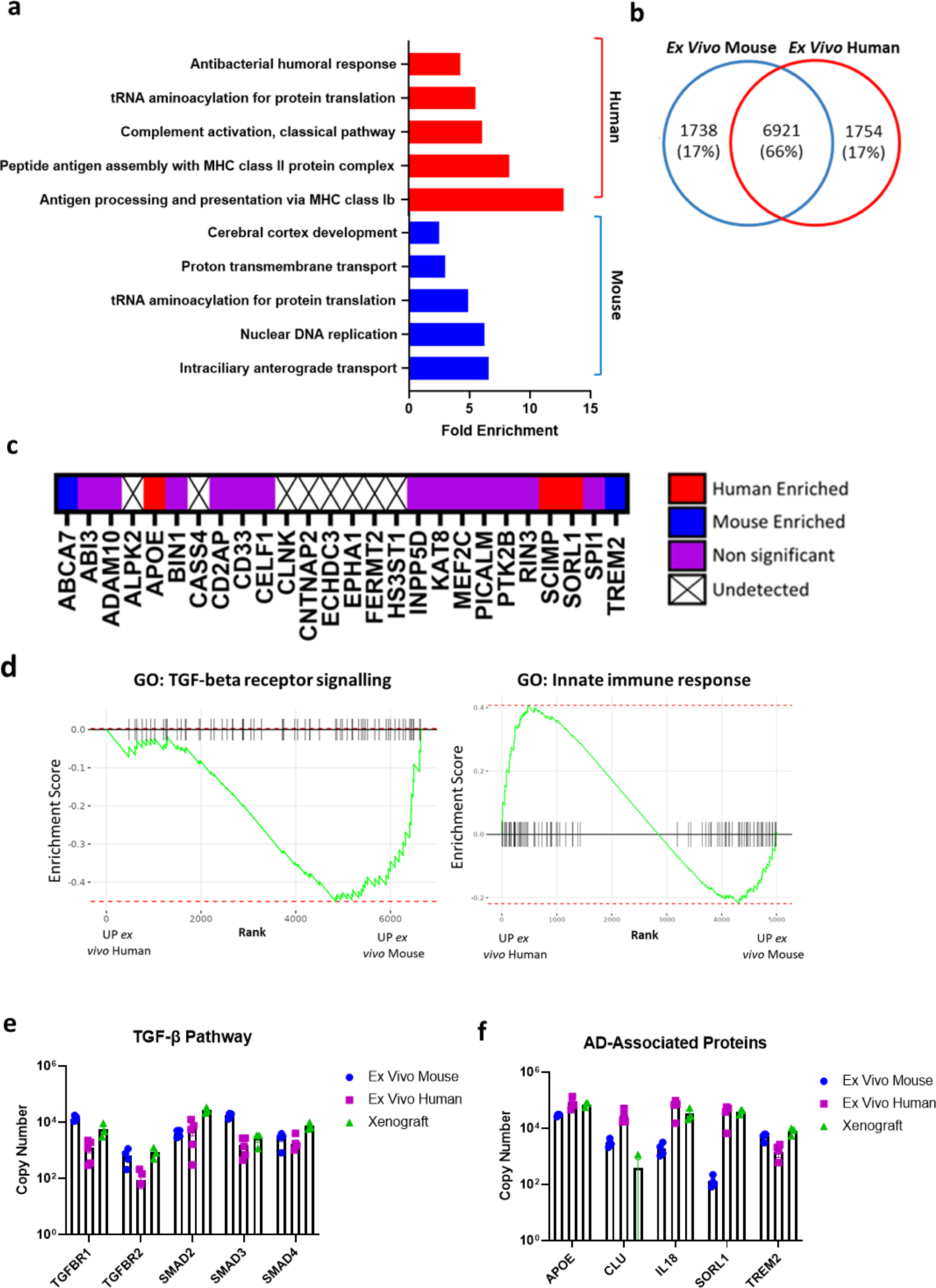
**a)** Go terms for proteins found exclusively in human (red) or mouse (blue) microglia, including species-specific orthologs. **b)** Venn diagram of proteins detected in both mouse and human microglia. **c)** Comparison of 26 Alzheimer’s Disease-associated proteins with shared mouse and human orthologs and their expression in *ex vivo* human and mouse microglia. Red = significantly enriched (p-adj<0.05), or only expressed in human microglia, blue = significantly enriched (p-adj<0.05) or only expressed in mouse microglia, purple = expressed in both, no significant differences in expression, white = not found in either. **d)** GSEA highlighting enrichment of TGF-β receptor signalling in *ex vivo* mouse microglia compared to *ex vivo* human microglia (left) and enrichment of proteins associated with innate immune responses in *ex vivo* human microglia compared to *ex vivo* mouse microglia. **e)** Copy number expression of proteins associated with TGF-β signalling in *ex vivo* mouse (n=4), *ex vivo* human (n=5) and xenograft microglia (n=3) ±SEM. **f)** Copy number expression of proteins associated with Alzheimer’s Disease (AD) in *ex vivo* mouse (n=4), *ex vivo* human (n=5) and xenograft microglia (n=3) ±SEM.

**Supplemental Figure 5.**
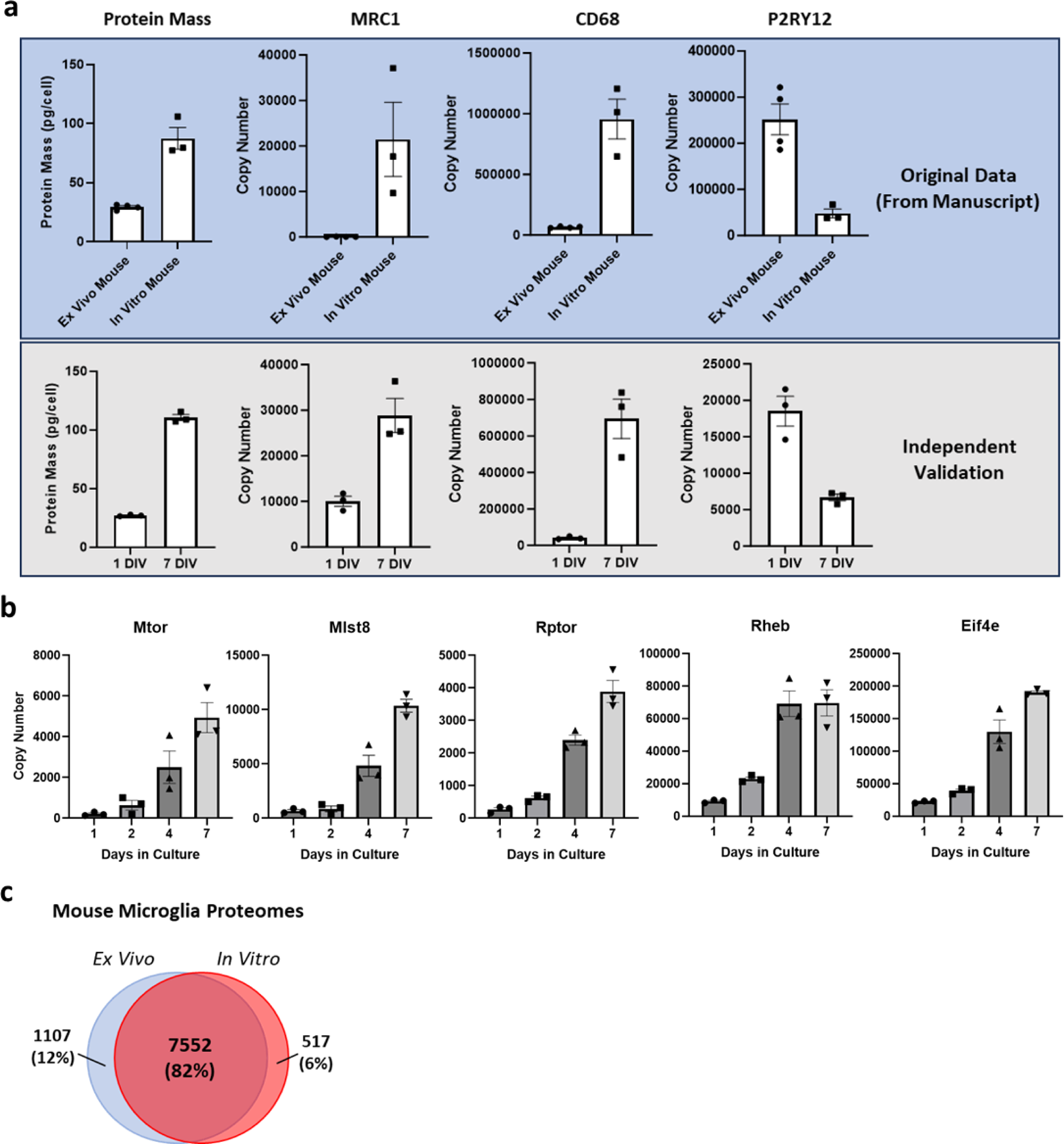
**a)** Comparing the total protein mass and the expression profile of key inflammation and homeostatic microglia proteins in *ex vivo* and *in vitro* microglia in the original dataset and the validation dataset. Proteomics differences are conserved across experiments. **b)** Copy number expression of MTORC1 proteins (MTOR, MLST8, RAPTOR), RHEB and EIF4E in *in vitro* mouse microglia over time in culture (n=3) ±SEM. **c**) Venn diagram representing the similarities and differences in proteins identified in *ex vivo* and *in vitro* mouse microglia proteomes. Of all proteins detected in *ex vivo* and *in vitro* mouse proteomes, 7552 (82%) were found in both, with 1107 (12%) only found in *ex vivo* mouse microglia proteomes and 517 (6%) only found in *in vitro* mouse microglia proteomes.

**Supplemental Figure 6.**
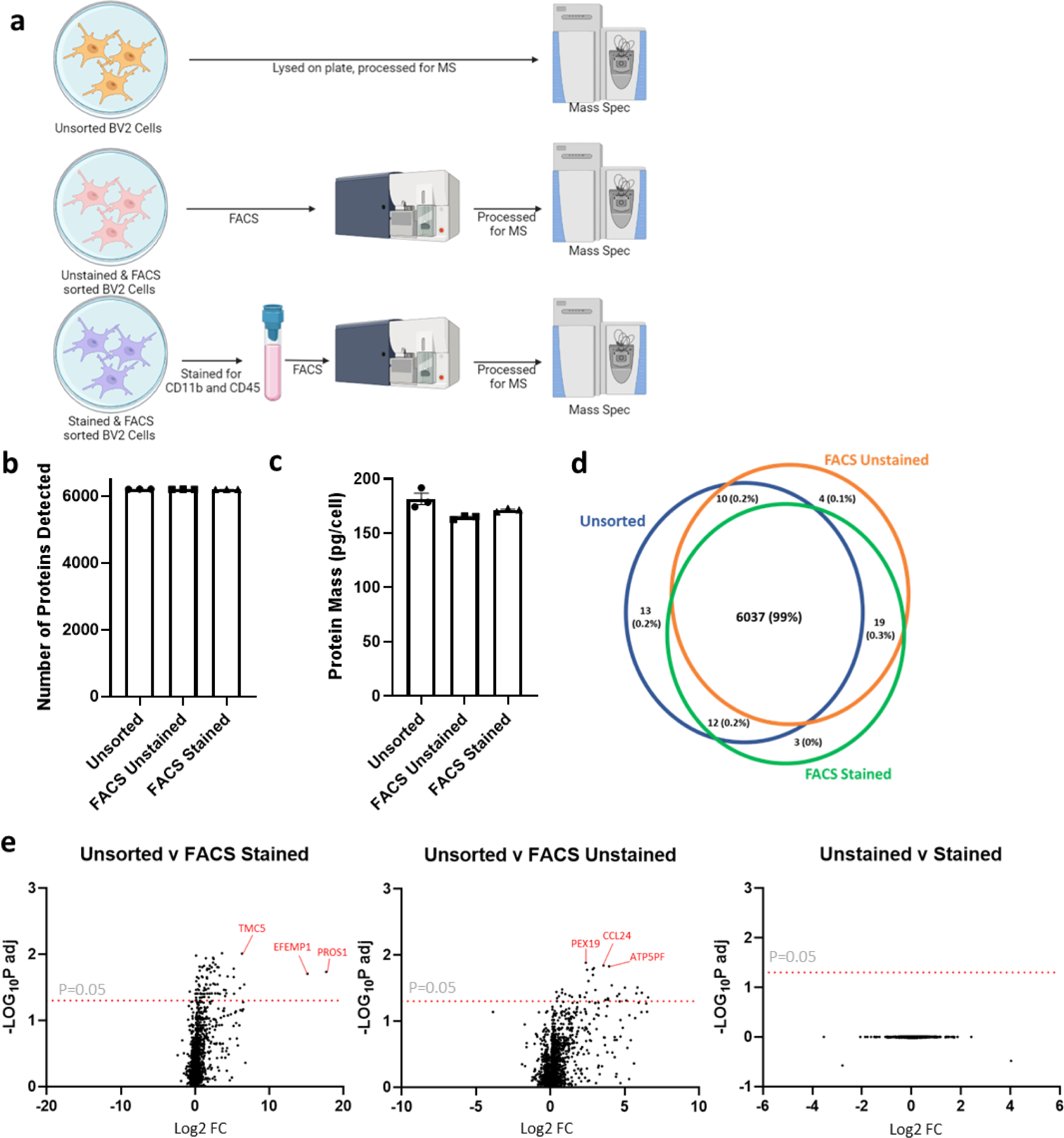
**a)** Schematic of experimental plan to investigate if sorting cells affects their proteome. Comparison of BV2 cells lysed directly on plate compared to processing and sorting by FACS either stained or unstained for CD11b and CD45, all processed for mass spectrometry. **b)** Number of proteins detected for unsorted or FACS sorted (stained and unstained)) BV2 cells ±SEM. N=3. **c)** Protein mass (pg per cell) of unsorted or FACS sorted (stained and unstained)) BV2 cells ±SEM. N=3. **d)** Venn diagram of all proteins identified in unsorted or FACS sorted (stained and unsorted) BV2 cells. **e)** Volcano plots of differentially expressed proteins between unsorted and FACS stained, unsorted v FACS unstained and FACS unstained v FACS stained BV2 cells. Data points above the red dotted line (p-adj=0.05) are statistically differentially expressed.

**Supplemental Figure 7.**
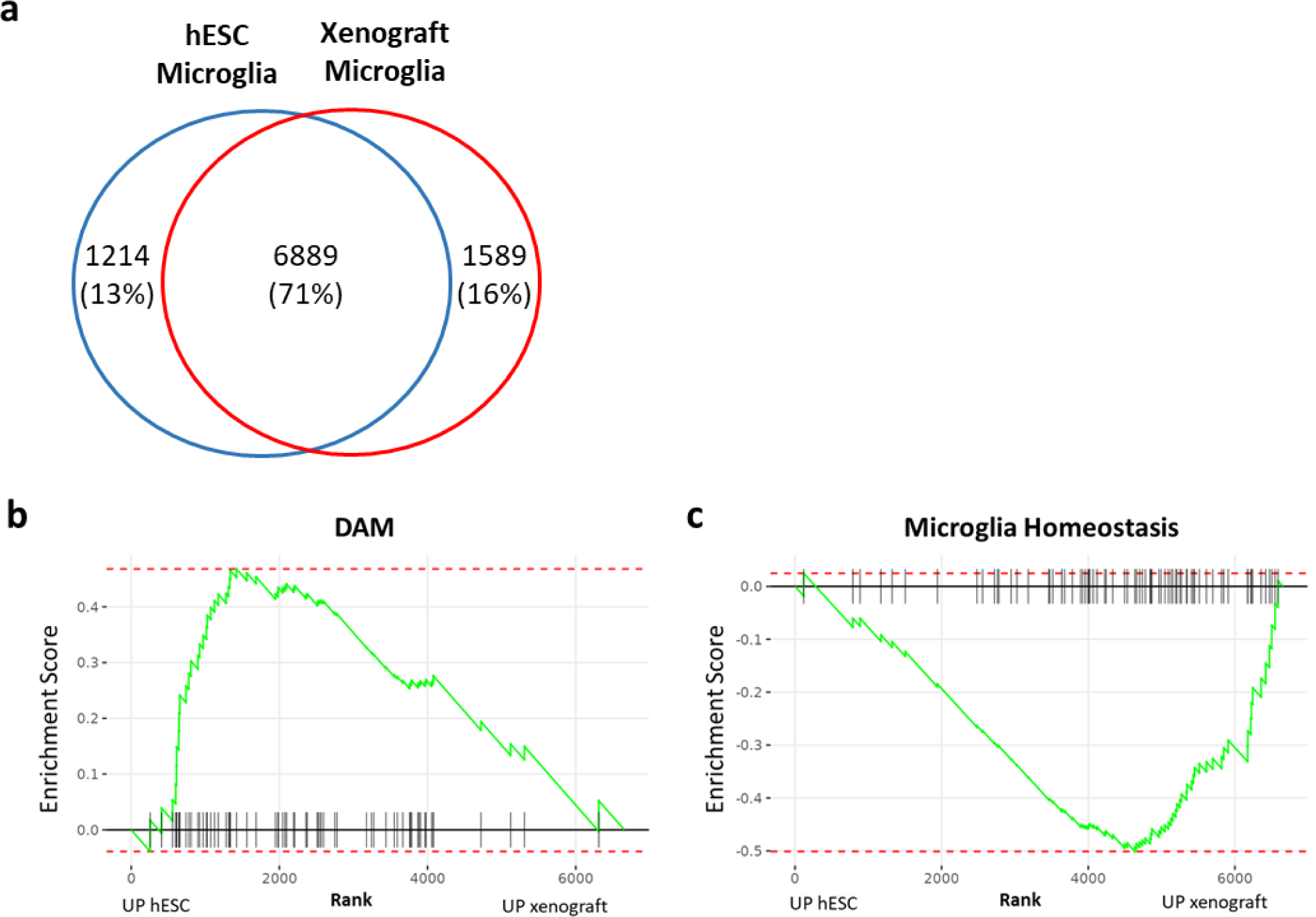
**a)** Venn diagram of proteins detected in *in vitro* hESC-microglia (blue) and xenograft microglia (red). **b)** GSEA of DAM proteins showing enrichment in *in vitro* hESC-microglia compared to xenograft. **c)** GSEA enrichment of Microglia Homeostasis proteins in xenograft microglia compared to *in vitro* hESC-microglia.

